# Oscillations in a Spatial Oncolytic Virus Model

**DOI:** 10.1101/2023.12.19.572433

**Authors:** Arwa Abdulla Baabdulla, Thomas Hillen

## Abstract

Virotherapy treatment is a new and promising target therapy that selectively attacks cancer cells without harming normal cells. Mathematical models of oncolytic viruses have shown predator-prey like oscillatory patterns as result of an underlying Hopf bifurcation. In a spatial context, these oscillations can lead to different spatio-temporal phenomena such as hollow-ring patterns, target patterns, and dispersed patterns. In this paper we continue the systematic analysis of these spatial oscillations and discuss their relevance in the clinical context. We consider a bifurcation analysis of a spatially explicit reaction-diffusion model to find the above mentioned spatio-temporal virus infection patterns. The desired pattern for tumor eradication is the hollow ring pattern and we find exact conditions for its occurrence. Moreover, we derive the minimal speed of travelling invasion waves for the cancer and for the oncolytic virus. Our numerical simulations in 2-D reveal complex spatial interactions of the virus infection and a new phenomenon of a periodic peak splitting. An effect that we cannot explain with our current methods.

## 1. Introduction

Cancer is a complex and devastating disease that requires extensive multidisciplinary research. Significant mathematical and computational modeling has been done to shed light on cancer formation and treatment. Many therapies have been developed over the last decades including radiotherapy [65; 52; 8], chemotherapy [86; 67; 69], surgery [49], immunotherapy [91] and virotherapy [71; 26; 37]. Compared to the conventional treatments that cause normal cell death as well as cancer cell death, new strategies have been developed to overcome this issue. Oncolytic virotherapy is considered one of the new promising target therapies. Oncolytic viruses selectively attack cancer cells and replicate rapidly inside them, thereby inducing tumor cell lysis. Hence, infecting the neighborhood cells and triggering the anti-cancer immune response. Moreover, virotherapy treatment can be used in combination with traditional treatments and with other target therapies such as immunotherapy and radiotherapy [84; 76; 83; 19; 38]. The mathematical modelling of oncolytic viraltherapy is at full swing (see the references below). In this paper we extend a spatially explicit model of Pooladvand et al. [68] to more fully explore the rich spatio-temporal dynamics of the oncolytic virus model. We obtain interesting spatial patterns and we discuss their relevance in the clinical context.

### 1.1. Oncolytical Virotherapy

Oncolytic virotherapy was inspired by clinical observations of tumor remissions after natural virus infections [45], prompting further investigation by clinicians and researchers [25; 74]. While a wide range of viruses have been used as oncolytic viruses in experiments such as adenovirus [30; 47; 33; 55], herpes simplex virus [72; 77], vaccinia virus [81], measles virus [7], reovirus [87; 62; 35; 59; 12] and vesicular stomatitis virus (VSV) [39], only two types have been licensed as anti-cancer treatments. These are adenovirus H101 for the treatment of head and neck cancer by Shanghai Sunway Biotech in 2005 [32], and herpes virus talimogene laherparepvec (T-VEC) for the treatment of advanced melanoma by U.S. Food and Drug Administration and European Medicines Agency [36] in 2015 and 2016, respectively.

Tumor control or tumor extinction is the main goal in cancer treatment. However, even with the advances in the pre-clinical and clinical trial results, analyzing and understanding the dynamics of tumor-virus interactions has proven to be difficult by experimentation alone. There are many challenges that virotherapy treatment faces to achieve the best results [44; 93]. For example, what is the balance of antiviral versus anti-tumor immunity? How can combination schedules and doses be optimized to achieve maximum anti-tumor efficacy? What is the best protocol design based on tumor type and oncolytic virus type? How does the spatial distribution of tumor and virus impact viral propagation and tumor control? Answering these questions is time consuming, expensive, and sometimes hard to address experimentally. Mathematical modelling can accelerate the progress of oncolytic virotherapy research by identifying crucial parameters, generating new hypotheses to be tested experimentally, predicting therapeutic outcomes in silico, and optimizing combined treatments. We note that the majority of mathematical modeling has focused on temporal dynamics of tumor-virus or tumor-virus-immune interaction because of the availability of temporal data. Some of these models are based on ordinary differential equations [43; 5; 23; 22; 70; 64; 82; 42; 79], while other models are based on delay differential equations [13; 15; 89; 54; 90]. Recent advances in intravital imaging and viral plaque analysis have been used to provide data of spatial spread of tumors and viruses [46; 80]. Many experimental studies [46; 80; 92; 28] assert the role of the spatial distribution on viral proliferation and spread within the host cells with the use of imaging and viral plaque analysis. Therefore, the influence of the spatial distribution on the tumor-immune-virus interaction cannot be ignored. Thus, several spatio-temporal models have been developed to understand the spatial dynamics of oncolytic viruses in cancer treatment [92; 10; 28; 24; 2; 66; 3; 68; 57; 56; 61; 78].

### 1.2. Modelling Oncolytic Virotherapy

To understand the principles that govern the virus spread through a target tumor cell population, Wodarz et al. [92] constructed an agent-based model based on in vitro experiments of adenovirus in a 2D setting of human embryonic kidney cells. Wodarz et al. [92] considered a spatially restricted domain with the assumption that the free viruses are in quasi steady state. The experimental results showed three spatial patterns: “*hollow ring structure* ”, “*filled ring structure*”, and “*dispersed patterns*”. The computational model results indicated that for successful treatment, the hollow ring structure is the best pattern since it was associated with either target cell extinction or a low level persistence of target cells.

A continuum version of Wodarz et al. [92] model in radial symmetry has been derived by Rioja et al. [15]. Rioja et al. [15] assumed the free viruses are not in the quasi steady state, therefore, the virus dynamics were expressed explicitly. Also, Rioja et al. [15] assumed the infected cells do not move and that the susceptible tumor cells and the free virus both decrease due to infection. As Rioja et al.’s model assumed radial symmetry, their analysis was restricted to radially symmetric patterns. The corresponding kinetic ODE system of Rioja et al. [15] model had been studied in [82].

Pooladvand et al. [68] explored a three-dimensional spherical symmetric model to investigate the dynamics of adenovirus in a spherical glioblastoma. They assumed the susceptible cells, infected cells and the virus particles can move. Furthermore, Pooladvand et al. [68] considered the loss of the free viruses due to the infection of susceptible and infected cells. Pooladvand and her colleagues [68] focused on the effectiveness of the infectivity parameter in the treatment outcome, where the virus was injected in the center of the tumor mass. Results showed enhancing the infectivity does not lead to a full eradication of the tumor. These results are consistent with the experimental results, suggesting that virotherapy treatment alone is often not good enough.

Bhatt et al. [6], investigated the reasons behind the virotherapy failures and how to improve the efficacy of the treatment based on the tumor cells sensitivity to the viral infection. To answer this question, Bhatt and his colleagues [6] used an immersed boundary method and a 2-D Voronoi and 3-D spatial interactions model. They find three main reasons for treatment failure: a high death rate of infected cells, leading to faster viral clearance; selection of virus-resistant cancer cells; and a too low viral spread rate.

Furthermore, Morselli et al. in [61] represented a stochastic agent-based model to study the impact of spatial constrains and the tumors microenvironment on the viral spread in solid tumors. Morselli et al. [61] considered two kinds of viral movements: undirected random movements and pressure driven movements and how this choice affects the virotherapy outcome. Their 2-D patterns from the agent-based model simulations were consistent with the patterns shown previously in Wodarz et al. [92] and Kim et al. [48].

To understand the role of immune system in virotherapy, several models had been developed in the literature [23; 22; 79; 78] and the references therein. One of the pioneer papers is the study of Storey et al. [79]. They develop an ODE framework model to investigate the role of innate and adaptive immune responses to virotherapy treatment of glioblastoma multiforme with Herpes Simplex Virus. The innate immune response has two functions in their model. On the one hand, it is a first responder, which attacks virus infected cells, and on the other hand, it stimulates the response of the adaptive immune system (T-cells). The authors find that under virotherapy alone, the virus clearance effect of the innate immune system dominates, the virus is removed quickly and the effect of the adaptive immune response is small. In an effort to boost the adaptive immune response, they consider a combination of virotherapy and immunotherapy in the form of a PD-1/PD-L1 inhibitor. The PD-1/PD-L1 pathway has been identified as a central mechanism that allows cancer cells to silence effector T-cells. If this pathway is inhibited, then the adaptive immune response stays active much longer and has a stronger effect on the cancer cells. Storey et al [79] show clearly a benefit from a combination therapy. In Storey and Jackson [78] they extend the model as a spatially explicit agent-based model. They show that to have an effective treatment, the viral dose should be injected in the location of highest tumor density rather than the tumor center.

Unfortunately, most of the spatial models mentioned above are computationally intensive, and further investigation into their pattern forming abilities is limited. In this paper, we focus on analyzing the viral infection pattern in cancerous tissue using a reaction diffusion formulation. This, will help us to find the best viral spread pattern that can lead to tumor eradication or at least keep the tumor under control. Furthermore, as cancer is spatially heterogeneous, considering the spatial effect will lead to a more realistic model and provide useful information leading to a better understanding of the tumor-immune-virus dynamics.

In this paper, we start from the detailed work of Pooladvand et al. [68]. They derived a radially symmetric model in 3-D for the interaction of cancer cells, infected cancer cells and the virus. A detailed bifurcation analysis showed the transition to a coexistent state as well as transition to oscillations via a Hopf bifurcation. These oscillations confirm previously reported oscillations in experiments by Kim [47] and Dingli [20]. Here we extend the model in various ways. We remove the assumption of radial symmetry and consider full 2-D or 3-D diffusion terms. We also remove the virus-loss term due to viral infection of cells, as this term is negligible. We analytically compute the invasion speeds of the model, and we use these to compare invasion profiles. Furthermore, we use the tumor control probability to quantify the likelihood that a tumor is eradicated. We find that in the oscillatory state, the tumor mass gets so small that it can be considered eradicated. Consequently, only the first oscillation period seems to be relevant and subsequent oscillations might be artificial. We also allow for general initial viral inoculation patterns (not necessarily centered at 0), which does have a significant effect on the viral spread patterns. Finally, as radial symmetry is not assumed, we find generic scenarios for very complex spaio-temporal patterns as they are known for spatially coupled oscillators in excitable systems [50; 4; 29; 73].

### 1.3. Outline

The paper is organized as follows: In the next Section 2 we introduce the mathematical model. We extend Pooladvand’s reaction diffusion model and perform a non-dimensionalization to reduce the number of parameters. In Section 3 we first consider the ODE-version of our model. We find the steady states, their stability and identify the two bifurcations that are relevant here, a transcritical bifurcation and a Hopf bifurcation. We then establish the minimal invasion speed for the one-dimensional PDE version of the model. In Section 4 we perform a detailed numerical exploration in 1D and 2D. We first identify the base values of the model parameters, guided by the work of Pooladvand [68]. We then compute the tumor control probability for the ODE simulations, to get an idea about treatment outcomes. For the 1D model we perform simulations that confirm the theoretical wave speed from Section 3. The simulations in 2D show interesting viral spread patterns that include coexistence steady states, hollow ring patterns as well as complex spatio-temporal interactions. We finish with a Conclusion in Section 5 where we put the results into perspective of mathematical modelling in general and cancer research in particular.

## 2. The Basic Assumptions and the Mathematical Model

In this paper, we will study an extension of Pooladvand et al. [68] reaction diffusion model on a rectangular or smooth domain Ω *⊂* R*^n^*:

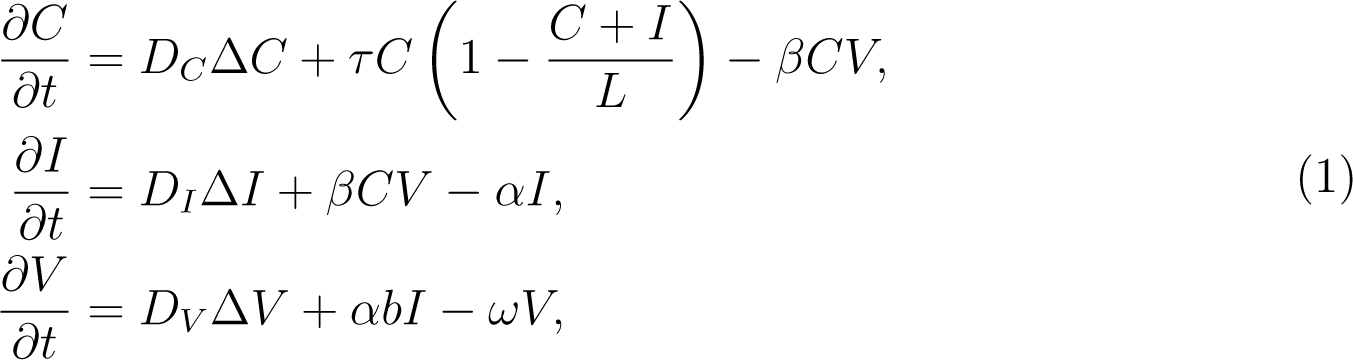

where the density of susceptible tumor cells, infected tumor cells, and free viruses is denoted by *C*(*x, t*), *I*(*x, t*), and *V* (*x, t*), respectively. The tumor growth rate is *τ* and *L* is the carrying capacity of tumor cells. The mass action term *βCV* describes the interaction between virus particles and the tumor cells. The infection rate is denoted by *β.* The death rate of infected cells is denoted by *α.* The burst size of the virus in infected cells is denoted by *b,* and *ω* is viral clearance rate. The diffusion coefficients of susceptible cells, infected cells, and viruses are denoted by *D_C_, D_I_, D_V_,* respectively. The symbol Δ denotes the Laplacian, i.e. the sum of all second order derivatives.

Compared to Pooladvand et al. [68] we relax the assumption of radial symmetry, and we removed the viral-loss term due to infection of cancer cells. We argue, as many other authors do as well, that the number of virions that are binding on a cancer cell is very small compared to the overall number of free virus particles. Hence this term can be ignored.

We consider system (3) with the initial conditions:

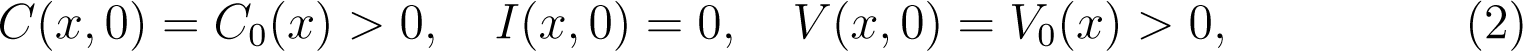

and homogenous Nuemann boundary conditions on *∂*Ω:

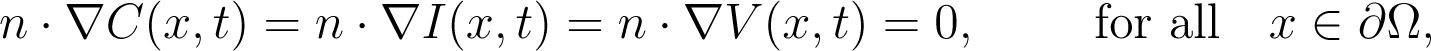

where *n* denotes an outward normal at *x* at *∂*Ω. In the case of a rectangular domain some points will have several outer normal vectors. In that case we require the above boundary conditions for all normal vectors. We use no-flux boundary conditions as is done in Pooladvand [68]. These boundary conditions are motivated by the experiments in a petri dish, which were studied in Wodarz [92]. Moreover, if applied to a tumor in some tissue, we consider the domain large enough such that boundary effects on the tumor are negligable.

To simplify our analysis, we apply non-dimensionalization by considering

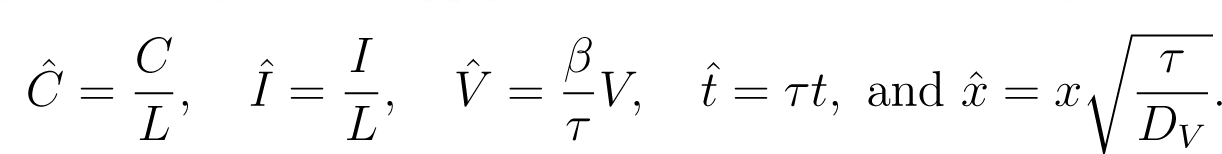

After dropping the hat, we get the non-dimensional system:

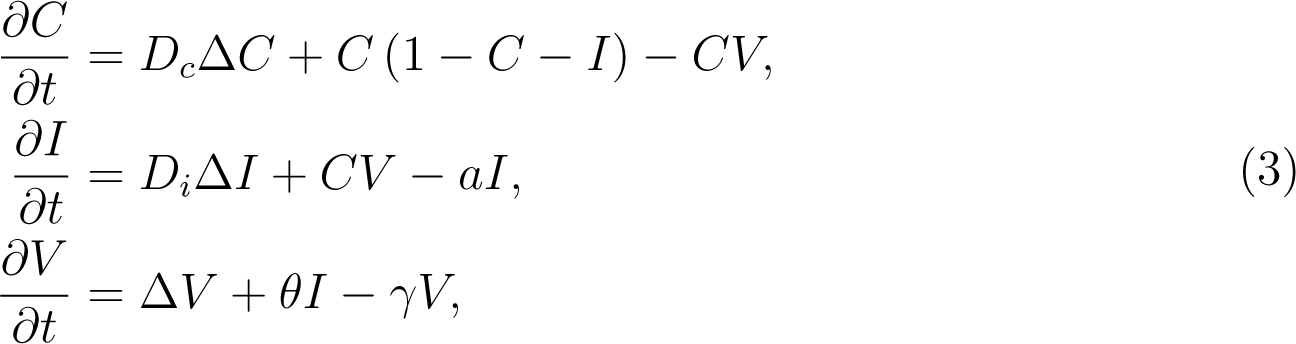

where

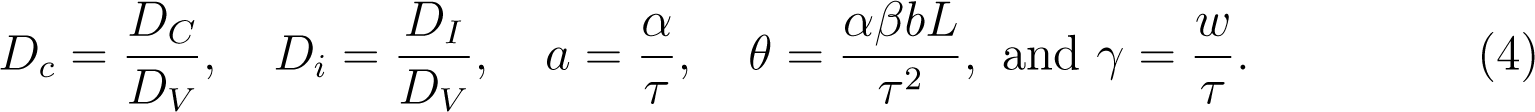

The effective viral production rate *θ* will be our bifurcation parameter in the next sections. It combines the death rate of infected cells *α*, the burst size *b*, the carrying capacity *L* and the cancer growth rate *τ*.

## 3. Analysis of the Model

### 3.1. Analysis of the Kinetic Part

We first consider the above model in the spatially homogeneous case, i.e. all spatial dependence is ignored. Then the system for *C*(*t*), *I*(*t*), *V* (*t*) becomes

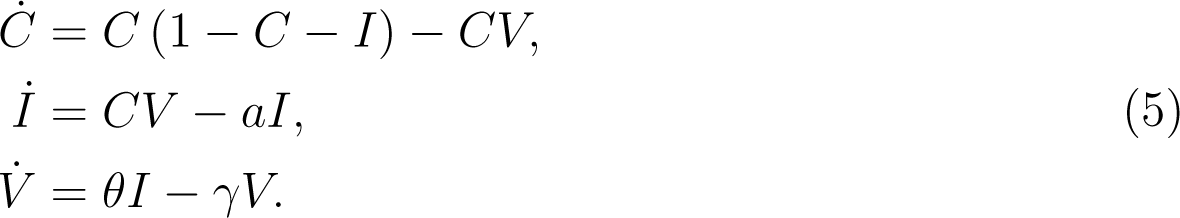

System (5) is of predator-prey type, and periodic orbits and Hopf bifurcations are standard properties of these models. Specifically [82; 68] have analysed similar models and found Hopf bifurcations. We also expect to find a Hopf bifurcation and oscillations. We summarize these results here, omitting some of the standard details.

The system (3) has three steady states which are the trivial steady state *E*_0_ = (0, 0, 0), the pure cancer state *E*_1_ = (1, 0, 0) and a coexistence steady state

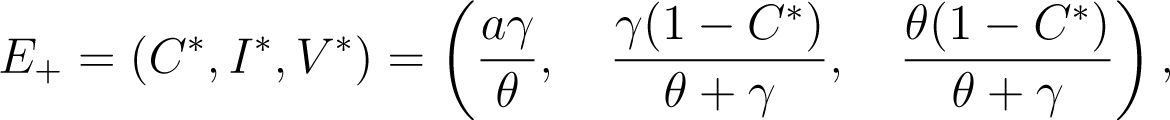

which exists in the positive quadrant for 0 *< C^∗^ <* 1. If *θ* is increased, the coexistence steady state arises through a transcritical bifurcation at

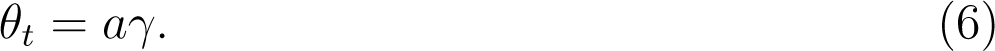

We compute the basic reproduction number *R*_0_ by applying the next generation matrix [18] with

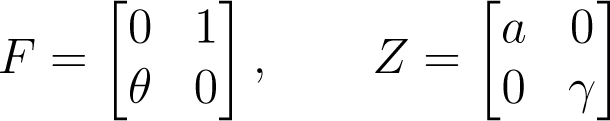

where *F* is the transmission matrix and *Z* is the transition matrix. Thus, 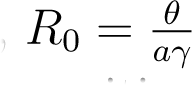 is the spectral radius of *FZ^−^*^1^. The coexistence steady state *E*_+_ exists in the positive quadrant iff *θ > aγ*, which is equivalents to *R*_0_ *>* 1.

It is straightforward to compute the stabilities of the steady states *E*_0_ and *E*_1_ and we omit some details here.

**Proposition 3.1.** Consider system (5).

1. *When R*_0_ 1, *we have two steady states E*_0_ *and E*_1_ *in the non-negative quadrant where E*_0_ *is a saddle point and E*_1_ *is locally asymptotically stable*.
2. *When R*_0_ *>* 1, *we have three steady states E*_0_, *E*_1_, *and E*_+_ *in the non-negative quadrant where E*_0_ *and E*_1_ *are saddle points*.
3. *The equilibrium point E*_+_ = (*C^∗^, I^∗^, V ^∗^*) *is locally asymptotically stable if θ > θ_t_ and κ*(*θ*) *>* 0, *where κ*(*θ*) = *−θ*^3^ + *mθ*^2^ + *γmθ* + *aγ*^3^, *and m* = (*a* + *γ*)^2^ + *a*(*γ* + 1).
4. *There exists a value θ_H_ > θ_t_ with κ*(*θ_H_*) = 0 *such that system (5) undergoes a Hopf bifurcation at θ_H_*.

**Proof.**

The third and fourth items are not so obvious and we give some details here. The Jacobian at *E*_+_ is

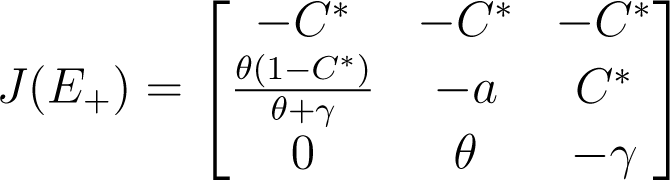

with the characteristic equation given by *λ*^3^ + *P*_2_*λ*^2^ + *P*_1_*λ* + *P*_0_ = 0, where

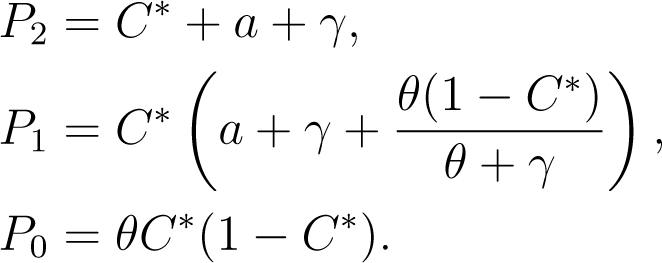

Based on Routh-Hurwitz criterion, the equilibrium point *E*_+_ = (*C^∗^, I^∗^, V ^∗^*) is locally asymptotically stable if *θ > θ_t_* and the following conditions hold:

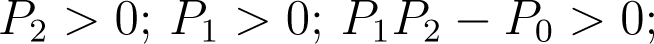

It is clear that *P*_2_, *P*_1_, and *P*_0_, are positive when 0 *< C^∗^ <* 1. It remains to show that *P*_2_*P*_1_ *− P*_0_ *>* 0. We find

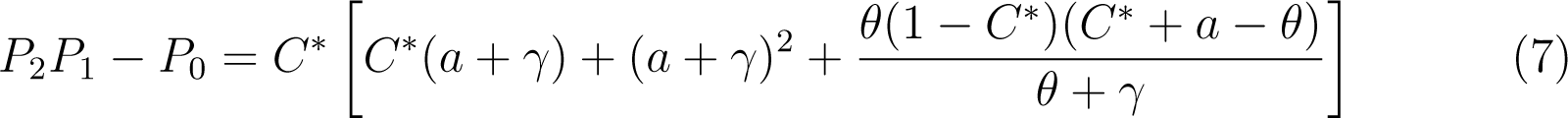

Substituting 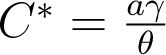 in equation (7), and after simplifications and re-arrangements, we get

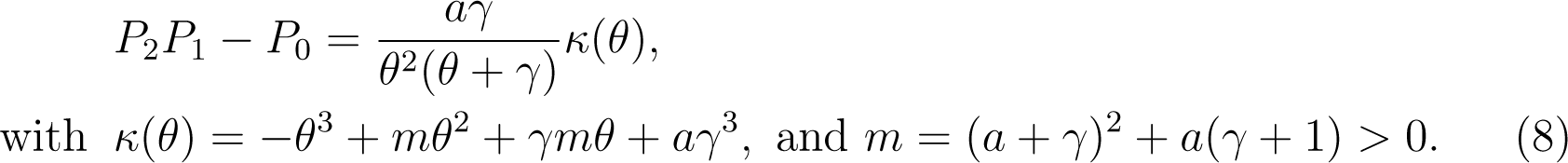

Since 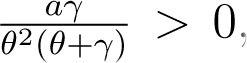, and *θ ̸*= 0, the stability of the coexistence steady state *E*_+_ is determined by the sign of *κ*(*θ*) after fixing the parameters *a* and *γ*.

To show the Hopf-bifurcation in item 4, we define

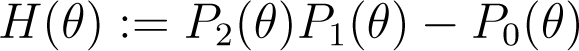

and use Liu’s criterion [51; 40], which says: Assume that *P*_0_ = 0. A Hopf bifurcation occurs at *θ* = *θ_H_* for the system (5) when the two following conditions are satisfied: 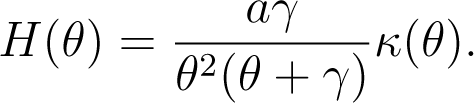.

From (8) we have

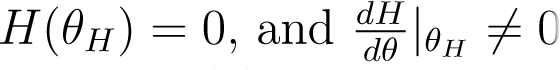

Thus, *H*(*θ*) = 0 iff *κ*(*θ*) = 0. Based on Descartes’ rule of signs, we have one positive real root and 2 or zero negative real roots. Since *θ* is a positive real number, there is a unique value of *θ* = *θ_H_,* where *H*(*θ_H_*) = 0. We also note that the leading order term in *H*(*θ*) is negative. This means for all *θ > θ_H_* we have *κ*(*θ*) *<* 0, which implies *θ_t_ < θ_H_*.

To obtain the direction of bifurcation we differentiate *H*(*θ*) as

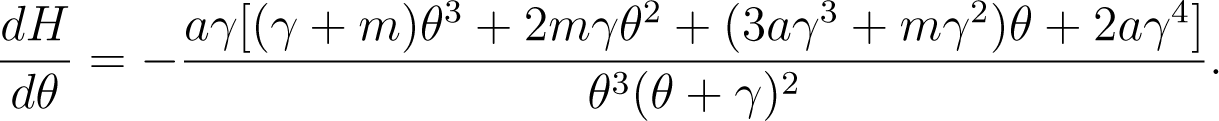

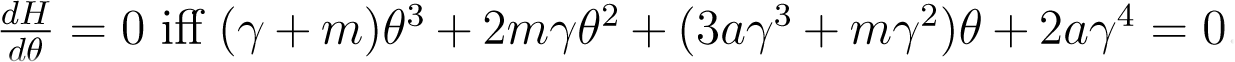. Again, using Descartes’ rule of signs, we find all the coefficients of *θ*’s are positive. Therefore, we have a no positive real roots. Thus, 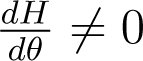 for any *θ.* Satisfying the Hopf bifurcation conditions.

We plot *κ*(*θ*) and the two bifurcation points *θ_t_* and *θ_H_* in Figure 1 with parameters from Table 2. In Figure 1 we also show some typical simulations of (5) with parameters from Table 2, for increasing values of *θ*. In this case *θ_t_* = 44.45 and *θ_H_* = 338.45. We see that as *θ* increases past *θ_t_* the coexistence steady state shows up (red dot), which becomes unstable to oscillations as we pass *θ_H_*.

**Figure 1:**
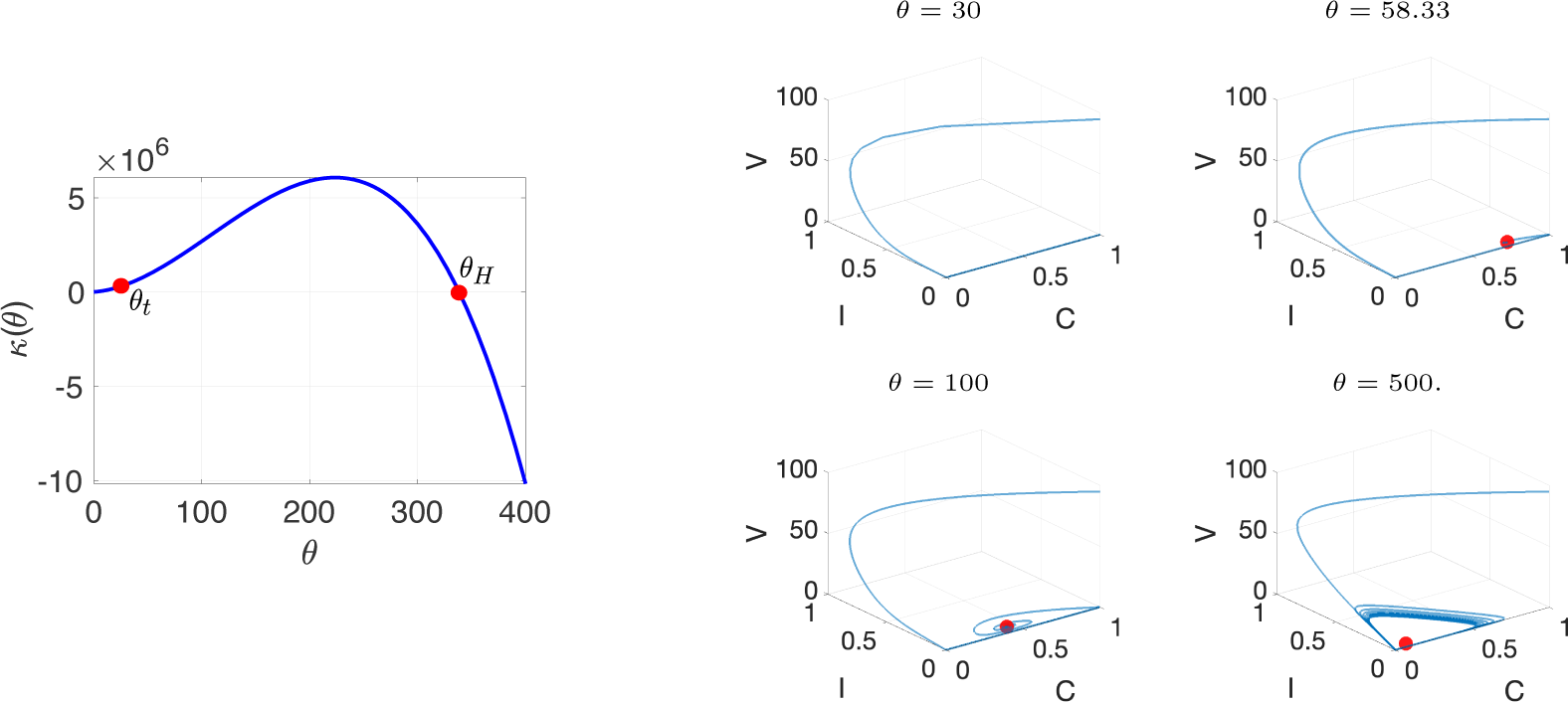
Left: Plot of the function *κ*(*θ*) from (8) for parameter values from Table 2. The two bifurcation points *θ_t_* = 44.45 for transcritical and *θ_H_* = 338.45 for Hopf bifurcation are indicated. Right: Numerical simulation of model (5) with different *θ* values with initial conditions C=1, I=0, and V=95. The coexistence steady state is indicated in red.

### 3.2. Wave Fronts of Glioma and Virus

In this section, we compute the invasion wave front of the spatial oncolytic virus model (3) in one spatial dimension by using a leading edge analysis. The domain is (*−∞, ∞*) and we consider first the tumor in isolation

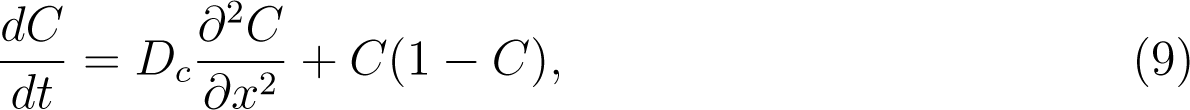

with boundary conditions

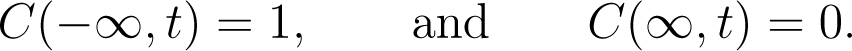

The standard traveling wave problem is to ask for a tumor invasion front into healthy tissue by considering a self-similar solution of the form *C*(*z*) with *z* = *x − c_c_t* that solves (9). Here *c_c_* ≥ 0 denotes the wave speed. We do not intend to recall all the details of the classical theory here, but we do need some details. Using the above wave ansatz, we derive a coupled ODE system for *C*(*z*) and *C^′^*(*z*) and our task is to find a heteroclinic connection from (1, 0) to (0, 0) in phase space. To find such a connection we linearize at (0, 0) and obtain a relationship between the wave speed *c_c_* and an exponential decay rate *λ_c_*, which is called the characteristic equation and can be written as

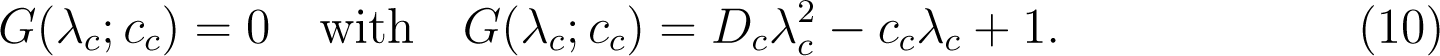

The minimum wave speed is the value such that *G* = 0 still has a real solution [16], i.e.

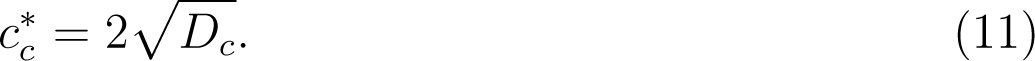

For later use we also linearize at the point (1, 0) in phase space. This point is a saddle point, i.e. it has a stable manifold along which solutions converge to (1, 0). The wave speed *c_c_* and the exponential decay rate on the stable manifold *λ_c_* are related as

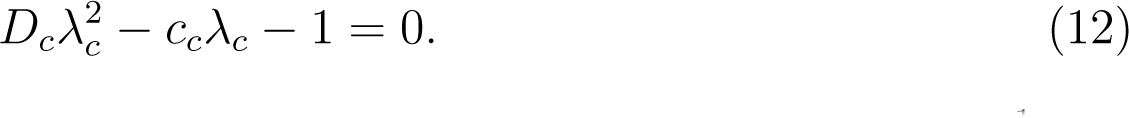

Note the opposite sign of the 1-term as compared to (10). For 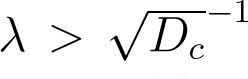 this equation has a solution for *c_c_*. However, in phase space for (*C, C^′^*) the unstable manifolds of (1, 0) do not connect to the other steady state (0, 0). In fact, going backwards in time they diverge to or and are not suitable solutions to our problem. Hence the condition (12) does not lead to a suitable invasion front.

To consider the invasion of the virus population into an established tumor, we linearize (3) in one dimension at the homogeneous steady state (1, 0, 0) and obtain

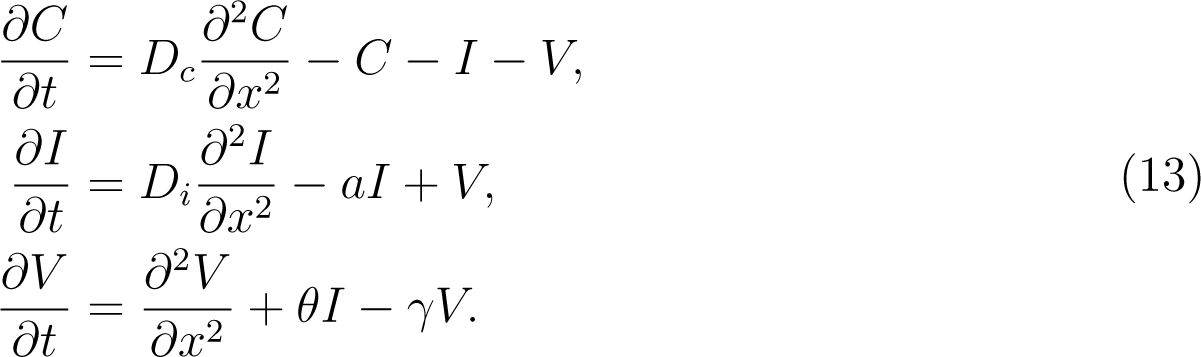

We now make an explicit ansatz of an exponentially decaying self-similar wave solution as

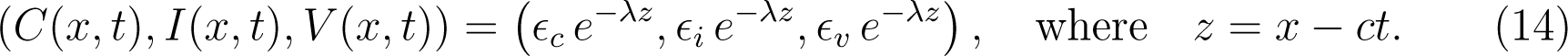

Here *c* denotes the wave speed and *λ* the exponential decay rate at the wave front. Substituting ansatz (14) into system (13), we get

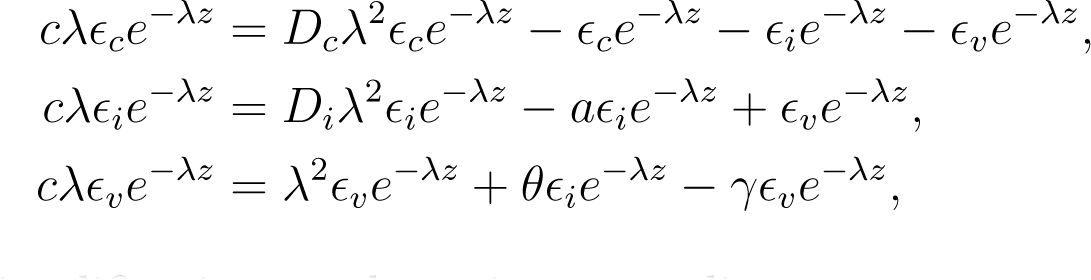

which after simplification can be written as a linear system

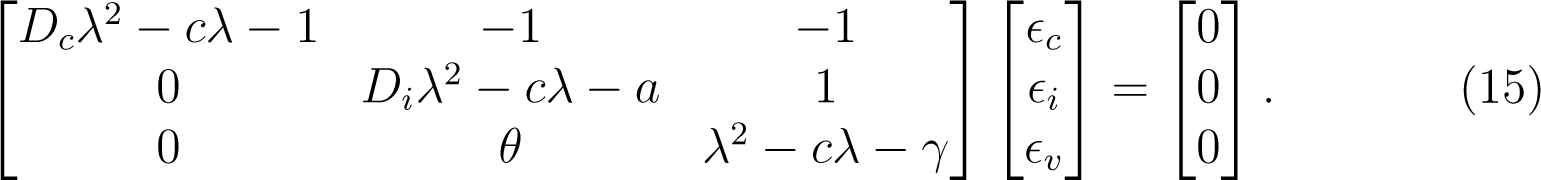

The characteristic equation of (15) is

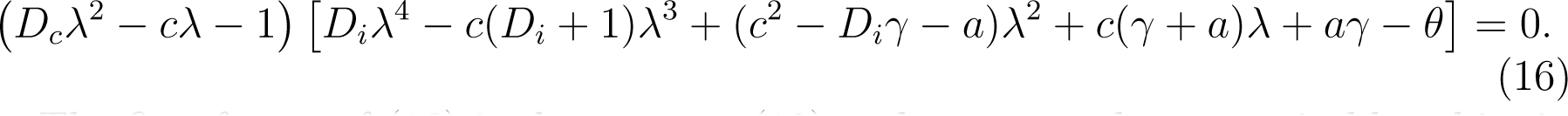

The first factor of (16) is the same as (12) and corresponds to unsuitable orbits in phase space. Hence we can focus on the second factor

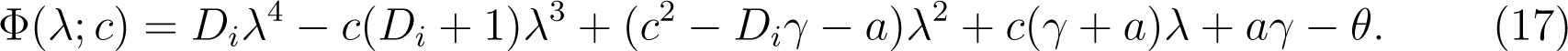

From Φ(*λ*; *c*) = 0 we can find the wave speed values *c*(*λ*) as functions of *λ.* To find the minimum wave speed of the virus, we use implicit differentiation

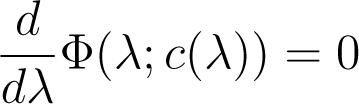

to find conditions for *c^′^*(*λ*) = 0. We have

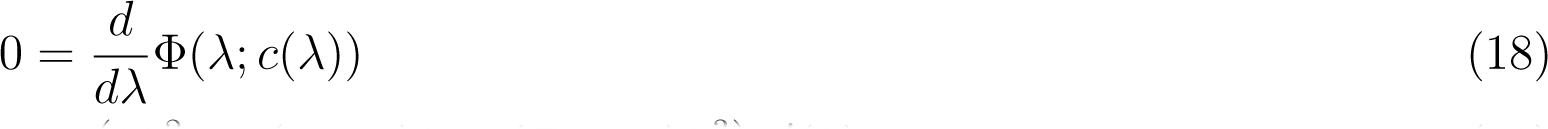

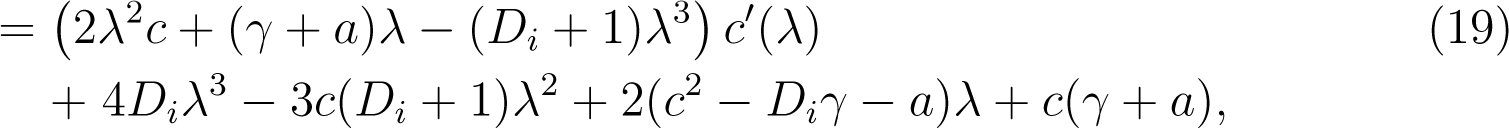

which can be solved for *c^′^* as

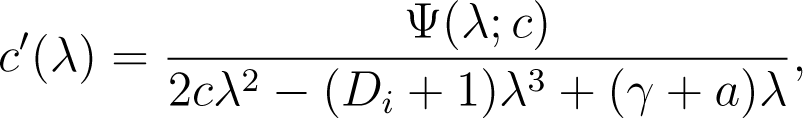

with

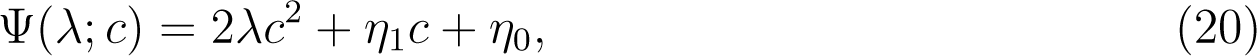

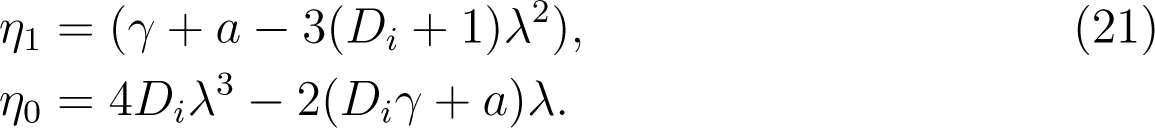

Thus, *c^′^*(*λ*) = 0, iff Ψ(*λ*; *c_v_*) = 0.

Hence the minimum wave speed *c^∗^* arises at the intersection of the two manifolds

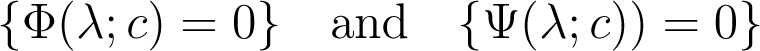

For the parameter values from Table 2 we plot these two curves in a (*λ, c*)-diagram in Figure 2. In that case we find a unique intersection at

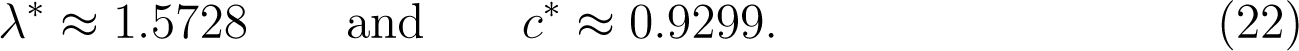

**Figure 2:**
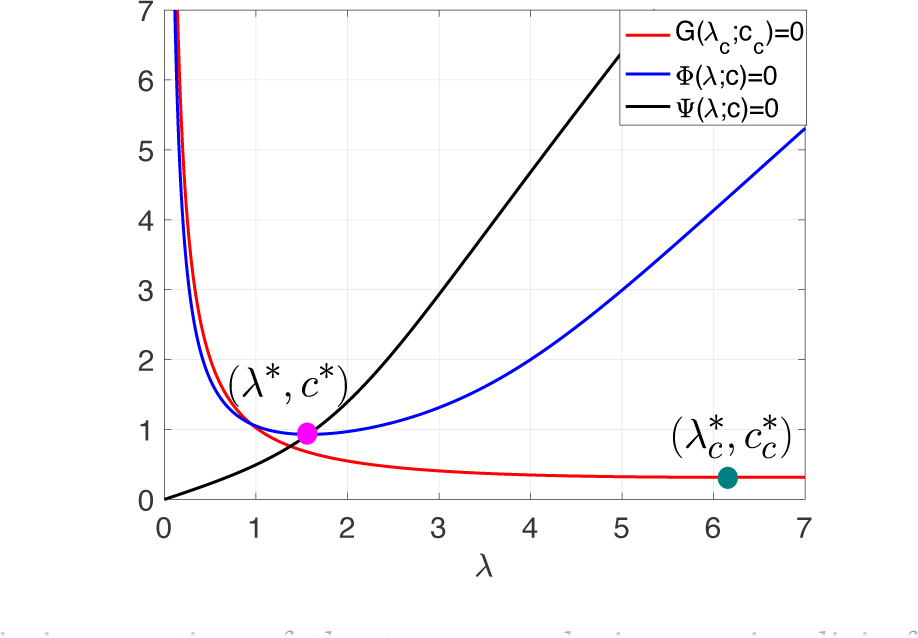
The characteristic equation of the tumor and virus as implicit function with respect to *λ* and *c.* The green circle is the minimum decay rate and corresponding minimum wave speed of tumor cells. The pink circle is the minimum decay rate and corresponding minimum wave speed of the virus particles. The parameters values are in Table 2.

This point is also indicated in Figure 2.

These values should be compared to the tumor invasion speed in isolation, which according to (11) is

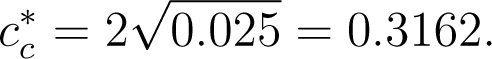

Moreover, we compute the minimal wave speed *c^∗^* and the decay rate *λ^∗^* for some of the parameter choices that we use later such as *θ* = 58.33, 100, 350, 500. The results are shown in Table 1. As *θ* increases, the decay rate *λ^∗^* becomes steeper, while the wave speed *c^∗^* increases.

**Table 1:** The decay rate *λ^∗^* and the corresponding wave speed *c^∗^* values across different effective viral production rates *θ*.

| $\theta$ | $\lambda^*$ | $c^*$ |
| --- | --- | --- |
| 58.33 | 1.5728 | 0.9299 |
| 100 | 2.4475 | 2.0227 |
| 350 | 4.0095 | 4.6958 |
| 500 | 4.4968 | 5.5400 |

## 4. Numerical Analysis

In this section, we present the numerical results for our model. First we explain the choice of model parameters in Section 4.1. Then, in Section 4.2 we illustrate the bifurcation path of the ODE model (5). The trivial steady state becomes unstable to a transcritical bifurcation at *θ_t_*, and shows oscillations after a Hopf bifurcation at *θ_H_*. We include a discussion of the tumor control probability (TCP) in Section 4.3 where we show that after one oscillation cycle, the tumor can be considered as removed. In Section 4.4 we consider the above model (3) in a one-dimensional situation and confirm the theoretical wave speeds that we found above. Simulations of a two-dimensional domain are shown in Section 4.5. We find the previously reported spread patterns of uniform spread, and hollow-ring patterns. In addition, for values of *θ* well beyond the Hopf point, we find unpredictable and chaotic patterns, which arise from spatial interactions of coupled oscillators via spiral waves and oscillating target patterns. Here we only touch on the rich variety of patterns that our system can generate and we leave a more complete analysis of this pattern regime for future work

### 4.1. Parameter Values

The parameters values that have been used in the simulations are presented in Table 2. These parameter values were used by Pooladvand et al. [68] in their study of adenovirus infections of a solid tumor. Based on Lodish [53], Pooladvand et al. [68] consider the carrying capacity of a solid tumor of radius 1 mm is about *L* = 10^6^ cells/mm^3^, which is also the initial density of uninfected tumor cells *C*_0_(*x*) in our model. Based on Kim’s et al [47] experiments on adenovirus in a glioblastoma U343 cell line, a tumor growth rate is estimated as *τ* = 0.3 per day, and a tumor diffusion coefficient as *D_C_* = *D_I_* = 0.006 mm^2^ per day. The initial condition for the amount of adenovirus virions is *V*_0_ = 1.9 10^10^ virions per mm^3^. For the viral diffusion coefficient, Pooladvand et al. [68] used the same value that has been employed in [66] which is *D_V_* = 0.24 mm^2^ per day. Since the infected cells lyse on average after 24 hours of injection [31], the death rate of infected cells is about *α* = 1 per day. The clearance rate of the virus has been estimated from [60] to be *ω* = 4 per day. The infection rate parameter *β* has been estimated based on Friedman’s et al. [28] paper to be *β* = 1.5 10*^−^*^9^ per (virus × day). Due to the importance of this parameter, Pooladvand et al. [68] varied the values of *β* from low to high, i.e *β* ∈ [10*^−^*^9^, 5 × 10*^−^*^9^] to investigate its impact on the treatment outcome. Shashkova et al. [75] estimated the adenovirus burst size *b* and found it varied between 1,000-100,000 virions per cell. Recently, Chen et al. in their paper [11] measured the burst size of adenovirus to be 3500 virions per cell, which is the value Pooladvand et al. [68] and we consider.

**Table 2:** Summary of CIV model parameters. All the parameters values used from reference [68].

| Parameter | Value | Units | Description |
| --- | --- | --- | --- |
| $C_0(x)$ | $10^6$ | cells per $\text{mm}^3$ | Initial density of uninfected cells |
| $V_0(x)$ | $1.9 \times 10^{10}$ | viruses per $\text{mm}^3$ | Initial density of virus particles |
| $D_C$ | 0.006 | $\text{mm}^2$ per day | Diffusion coefficient of susceptible cells |
| $D_I$ | 0.006 | $\text{mm}^2$ per day | Diffusion coefficient of infected cells |
| $D_V$ | 0.24 | $\text{mm}^2$ per day | Diffusion coefficient of viruses |
| $\tau$ | 0.3 | per day | Growth rate of tumor cells |
| $L$ | $10^6$ | cell per $\text{mm}^3$ | Carrying capacity of tumor cells |
| $\beta$ | $1.5 \times 10^{-9}$, variable | per (virus × day) | Infection rate of mass action case |
| $\alpha$ | 1 | per day | Death rate of infected cells |
| $b$ | 3500, variable | virus per cell | Burst size of viruses |
| $\omega$ | 4 | per day | Clearance rate of viruses |

In the virotherapy treatment the crucial parameters for successful treatment are the infection rate *β* and the viral burst size b. Therefore, in this paper, we will focus on the effective virus production rate parameter *θ* from (4), which includes these two parameters. Thus, using the values in Table 2, we have the base value of *θ* = 58.33. However, we will use *θ* as a bifurcation parameter in our study. Therefore, fixing the parameters L, *τ*, *α,* and *w,* as in Table 2, and varying *β* and b, we find the effective virus production rate *θ* is ranged between 11.1111-5555.55.

After rescaling (4), the parameter values for model (5) are given in Table 3.

**Table 3:** Base parameter values of the rescaled model (3) using the rescaling (4). These parameters are unit less.

| Rescaled Parameter | Meaning | Value |
| --- | --- | --- |
| $D_c$ | cancer diffusion coefficient | 0.025 |
| $D_i$ | infected cells diffusion | 0.025 |
| $a$ | infected death rate | 3.33 |
| $\gamma$ | virus decay rate | 13.33 |
| $\theta$ | effective virus production rate | [0,500] |

### 4.2. ODE Model

We start by studying the ODE system (5) with *θ* from (4) as a bifurcation parameter, where *θ* represents the effective virus production rate. In Figure 1 we show the dynamic behavior of model (5) with parameters from Table 3 with varying *θ*. The initial conditions are (*C*(0), *I*(0), *V* (0)) = (1, 0, 95) in all four simulations. In Figure 3 we show the cases of *θ* = 58.33, 100, 500 as functions of time. The transcritical bifurcation occurs at *θ_t_* = 44.45, and for *θ > θ_t_* we obtain a coexistence steady state. The Hopf bifurcation occurs are *θ_H_* = 338.45 and we obtain periodic orbits beyond that point. Oscillations arise already for *θ < θ_H_*.

**Figure 3:**
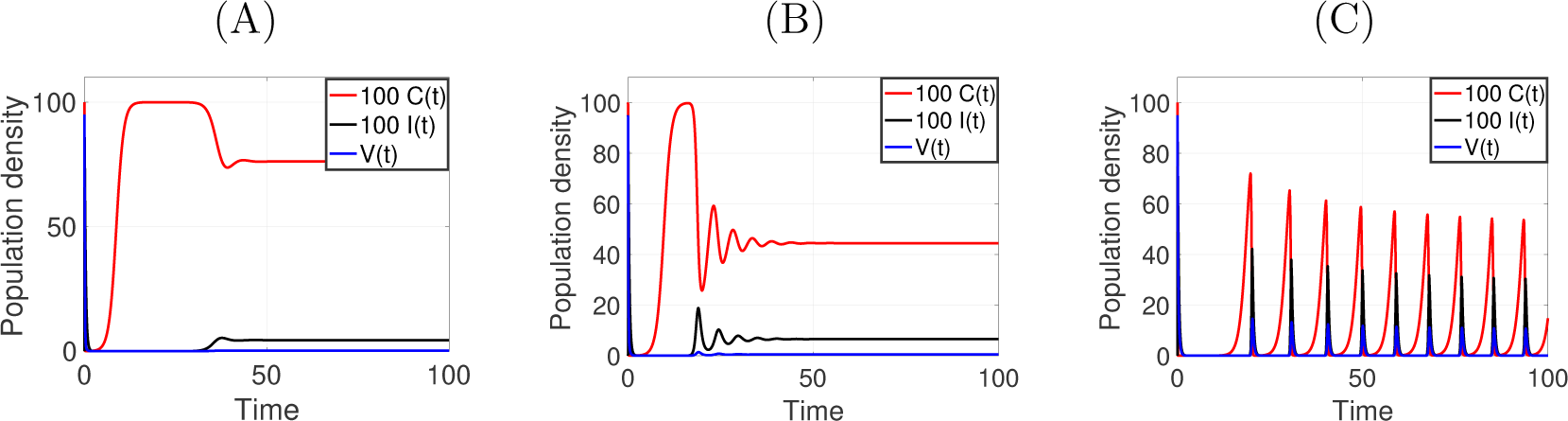
Numerical simulation with different *θ* values with initial conditions (1, 0, 95). Since, the initial virus particles at t=0 is 95, we scale up the cell densities by a factor of 100 such that these curves can be shown in the same coordinate system. **(A):** *θ* = 58.33, **(B):** *θ* = 100, and **(C):** *θ* = 500.

Note that in all cases the trivial steady state (0, 0, 0) is unstable, hence, at least mathematically, we can never fully eradicate the cancer. However, the periodic orbits get very close to (0, 0, 0). In fact, they get so close that we employ a stochastic view point using the tumor control probability in Section 4.3 below.

Before we do this, we briefly consider the role of the other model parameters.

In Figure 4 we vary the value of the virus decay rate *γ* from its base value of *γ* = 13.33 including the values *γ* = 6.67, 10, 16.67. We see a positive effect on cancer reduction for reduced virus decay rate. The bifurcation structure is essentially unchanged for the different *γ* values, but the oscillation frequencies change from low frequency for *γ* = 6.67 to high frequency for *γ* = 16.67.

**Figure 4:**
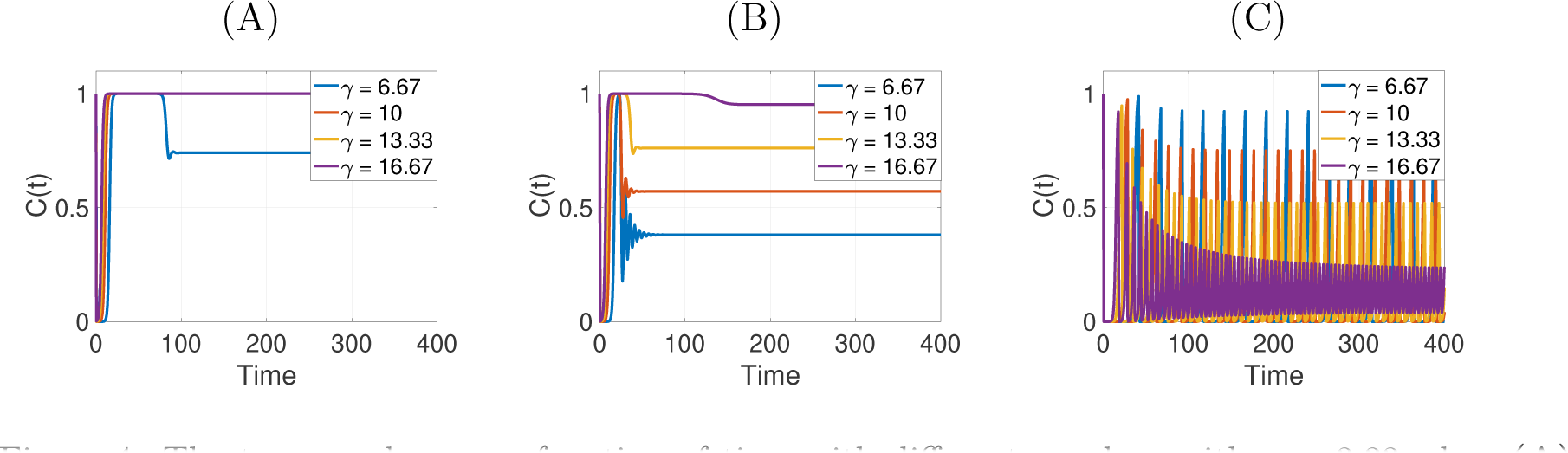
The tumor volumes as function of time with different *γ* values with *a* = 3.33 when **(A):** *θ* = 30, **(B):** *θ* = 58.33, **(C):** *θ* = 500.

In Figure 5 we vary the removal rate of infected cells *a* from its base value *a* = 3.33 and include *a* = 6.67, 10, 13.33. In this case we see that the bifurcations strongly depend on the value of *a*. For increased removal rate we lose the oscillations at *θ* = 500 and observe convergence to a coexistence steady state. Interestingly, increasing the removal rate of infected cells *a*, increases the value of the bifurcation point *θ_t_* = *aγ* of the transcritical bifurcation. Such a shift will change the treatment outcome as we can see in Figure 5. This observation is consistent with Bhatt et al. [6] results, where increasing the death rate of infected cells leads to treatment failure. Bhatt et al. [6] also identified the presence of resistant cancer cells as a reason for treatment failure. In our model we can address this through a reduced infection rate *β*. If *β* is reduced so is *θ*, i.e. moving the system to lower virus production rate such that the cancer grows easier.

**Figure 5:**
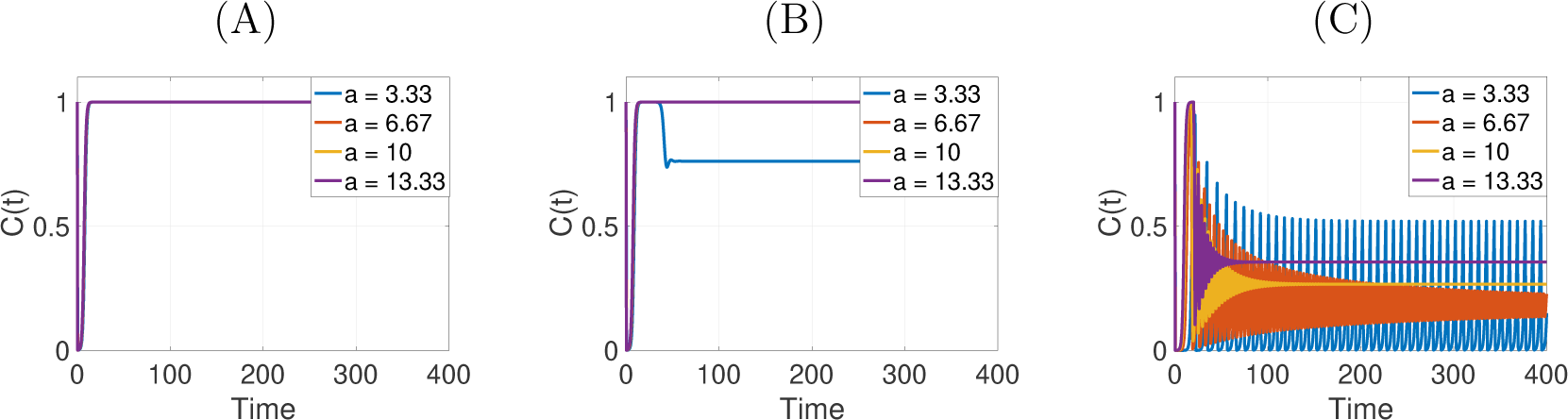
The tumor values with different *a* values with *γ* = 13.33 when **(A):** *θ* = 30, **(B):** *θ* = 58.33, **(C):** *θ* = 500.

### 4.3. Tumor Control Probability

Our main goal is tumor eradication. Since (0, 0, 0) is unstable for model (5) for *θ > θ_t_*, we will never be able to fully eradicate the cancer cells. However, we notice that at the oscillations described above, the tumor cell density gets very close to zero. In such cases we enter a region where stochastic events become important and a deterministic description via an ODE model might not be sufficient. Stochastic models for tumor eradication were developed for cancer radiation therapy [95; 94; 34; 65]. The tumor control probability (TCP) is a measure for the expected success of a given treatment. In Gong et al. [34] several formulations for the TCP were compared and it was found that the simplest version, the Poissonian TCP, is a very reasonable first order approximation. Hence here we use the Poissonian TCP. Let *C*_0_ denote the initial number of tumor cells, and *C*(*t*) those that survive at time *t*, then

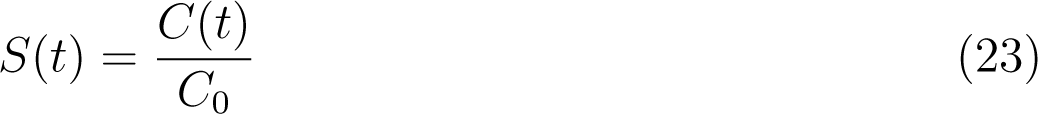

denotes the survival fraction and the Poissonian TCP is given as

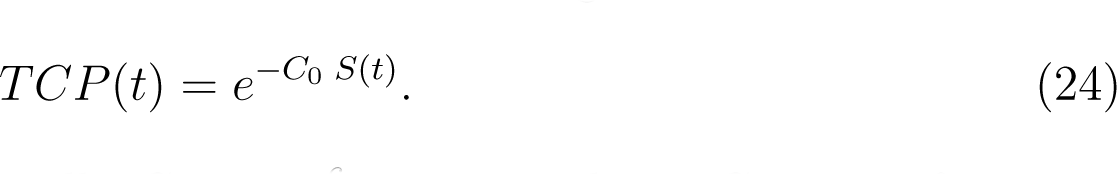

The initial number of tumor cells *C*_0_ = 10^6^ corresponds to *C*_0_ = 1 after non-dimensionalization.

We consider three different cases of *θ* = 58.33, 350, 500 and plot the TCP as a function of time in Figure 6. For the base case of *θ* = 58.33 the TCP first approaches 1 and then converges to a medium value of about 0.45. For *θ* = 350 and 500 we are beyond the Hopf point, and we see plateaus at values very close to TCP=1 for extended times right after virus infection treatment. This means that TCP=1 after the first cycle and the treatment is expected to be successful. The further oscillations are irrelevant for the treatment outcome, even though they exist mathematically. We come back to this issue when we discuss the two-dimensional simulations.

**Figure 6:**
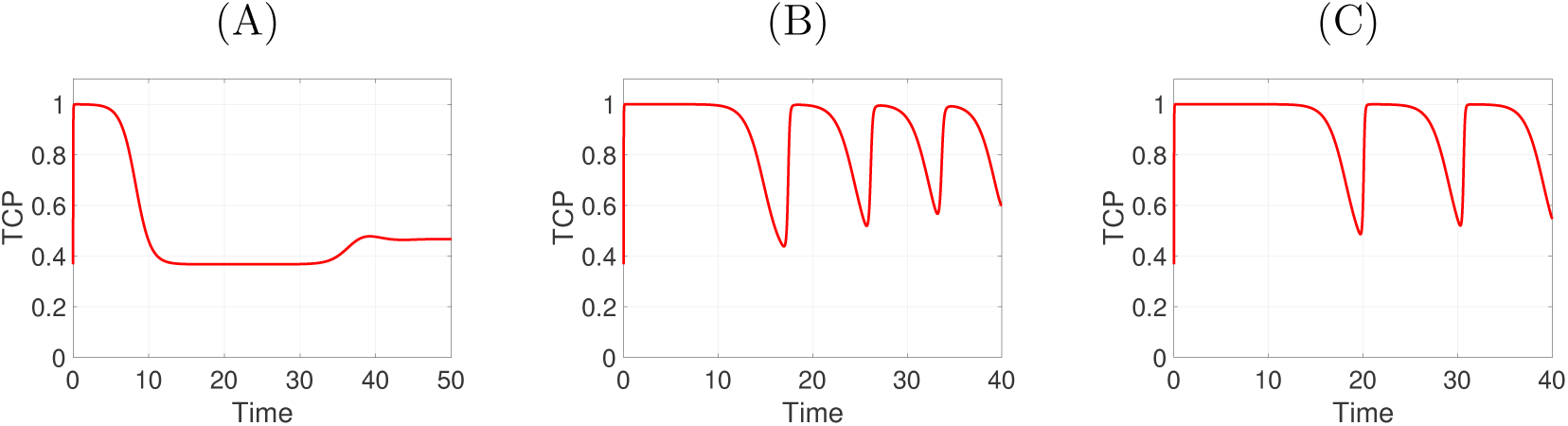
TCP values of tumor cells when *θ* in the Hopf bifurcation region at *a* = 3.33 and *γ* = 13.33. **(A):** *θ* = 58.33, **(B):** *θ* = 350, and **(C):** *θ* = 500.

### 4.4. PDE Model 1-D

As we understand the Hopf bifurcation of the ODE model well, it is interesting to see how the oscillations manifest themselves in the spatial context. It is known through the influential Kuramoto model [50] that spatially coupled oscillators can lead to all kind of dynamics such as ring patterns, target patterns, spirals and chaotic patterns [50; 73]. Those models have been applied with success in neurophysiological oscillations [9; 29], for example. We like to explore here if our model is also able to generate interesting patterns and evaluate them in the context of cancer treatment.

Starting in 1-D we consider an interval [0, 60] and run simulations up to time 60. In dimensional parameters this corresponds to an interval of length 54 mm and a time of up to 200 days. Initially the virus particles are injected in the center of the domain. We chose zero-flux boundary conditions. The simulations are seen in Figure 7. The cancer cell density *C*(*x, t*) in (A) and (D) is shown in colors of brown, where dark color corresponds to maximum tumor cell density. The infected cells *I*(*x, t*) in (B) and (E) are shown in shades of blue, with dark blue indicating low density of infected cell. In (C) and (F) we show the virus load in colors from blue (low) to red (high). In Figure 7 (A), (B) and (C), we chose *θ* = 58.33, i.e. *θ_t_ < θ < θ_H_*. The initial virus infection grows out from the centre of the domain and forms a travelling wave profile. After the wave has passed (around time 40) the system reaches a homogeneous coexistence steady state. We can use this simulation to estimate the travelling wave speed (slope of the ridges in (A), (B), (C)) as *c_v_* = 0.9428, while the theoretical wave speed (22) is *c^∗^* = 0.9299, which is a good match. On the other hand, when *θ* = 500, we are past the Hopf-point and we see oscillations in Figure 7 (D), (E) and (F). The oscillation synchronises spatially until it becomes a fully spatially homogeneous oscillation (long term dynamics not shown).

**Figure 7:**
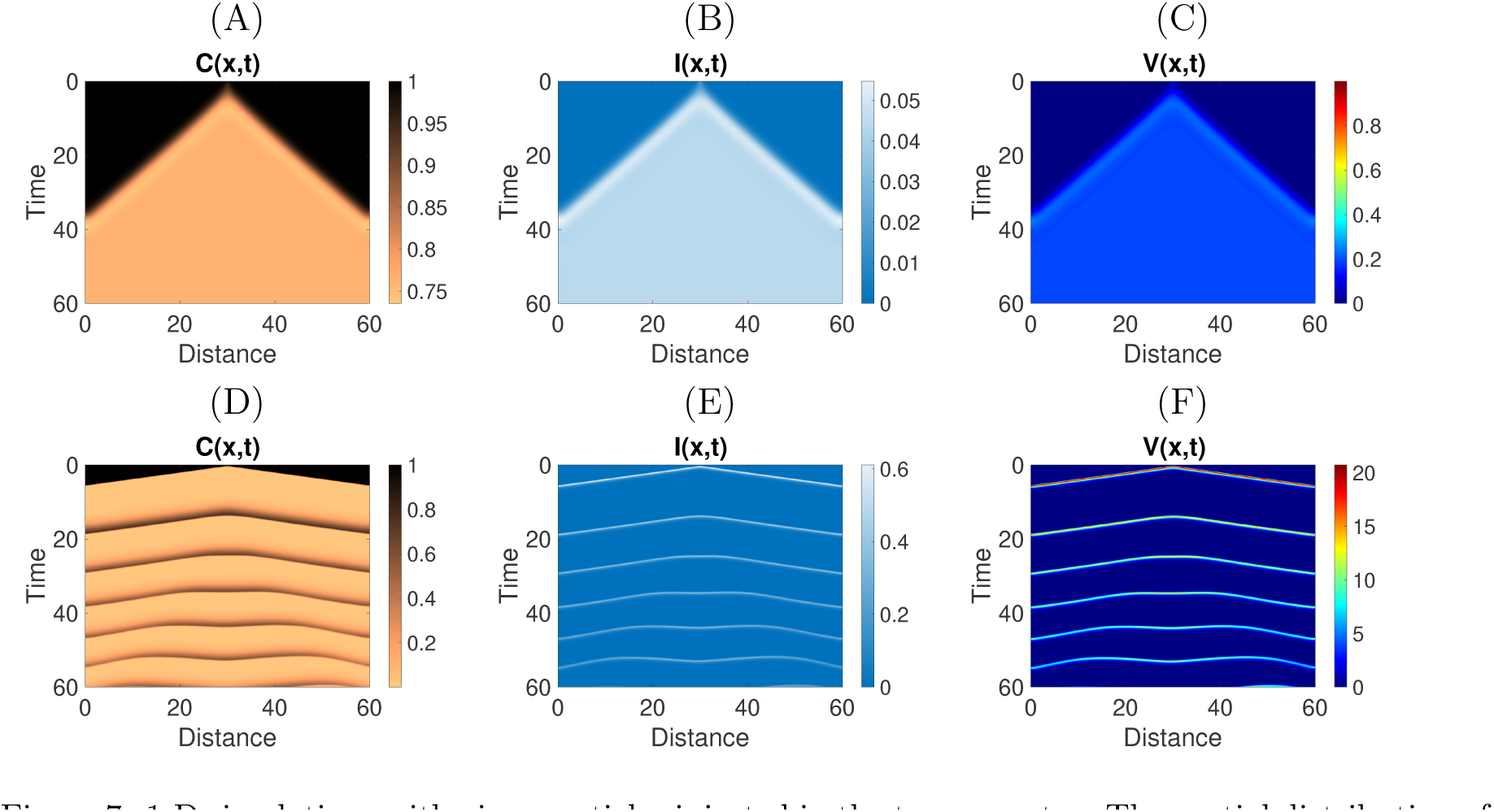
1-D simulations with virus particles injected in the tumor center. The spatial distribution of the susceptible cells, infected cells and the virus with different values of *θ* with *a* = 3.33 and *γ* = 13.33. θ = 58.33 for **(A)**, **(B)** and **(C)** and *θ* = 500 for **(D) (E)** and **(F)**.

We perform the same simulations with a different initial condition in Figure 8 (A), (B), (C). Here we randomly chose five inoculation points along the interval [0, 60]. For *θ* = 58.33 in (A) we see an initial interaction of the spread waves, which quickly find the coexistence steady state. For *θ* = 350 in (B) and *θ* = 500 in (C) we again see synchronized oscillations across the domain.

**Figure 8:**
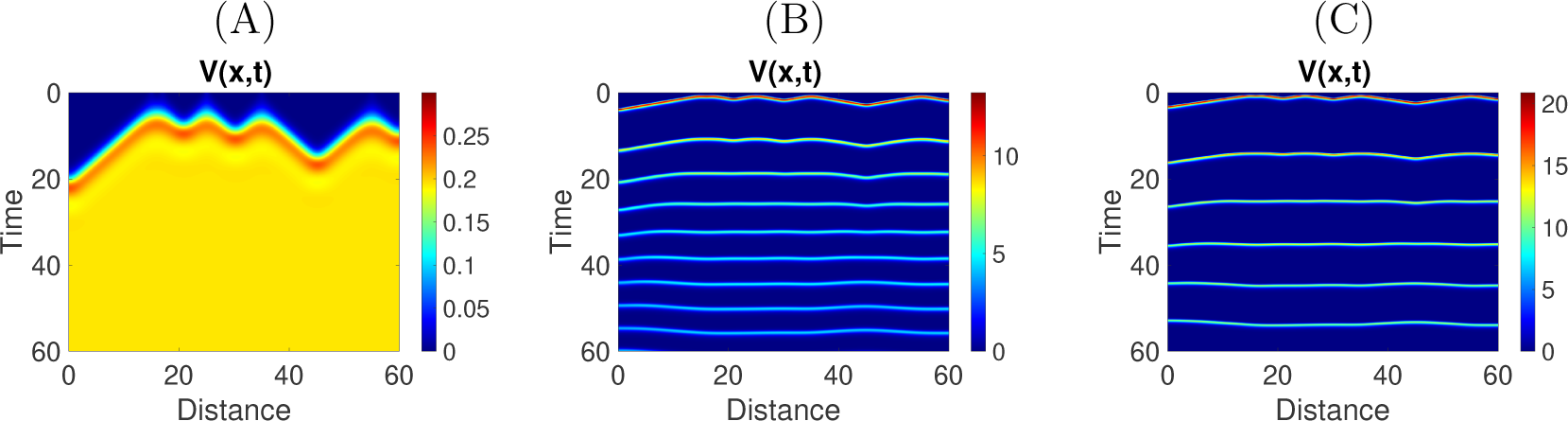
1-D simulations with virus particles injected in different regions of the tumor. **(A):** *θ* = 58.33, **(B):** *θ* = 350, and **(C):** *θ* = 500.

As in the ODE model the oscillations get very close to zero, indicating very low cancer and virus concentrations. Hence again, we employ the tumor control probability (24) to estimate the expected treatment success. In the spatial context we integrate over space to get the total surviving fraction of cancer cells. Then the TCP becomes

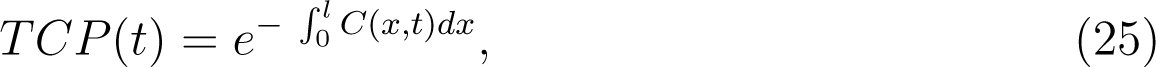

where we use the time point where the cancer concentration *C* is minimal. For the cases where we inject the oncolytic virus in the center of the domain we find that the maximal TCP value is 0 for *θ* = 58.33, 0.2598 for *θ* = 350, and 0.9753 for *θ* = 500. The TCP values were generally higher when the tumor was injected at five random locations with TCP of 0 for *θ* = 58.33, 0.8103 for *θ* = 350, and 0.9944 at *θ* = 500. This clearly shows that for the base value of *θ* = 58.33 treatment failure is expected. Only if *θ* is increased beyond the Hopf point we can expect good treatment outcomes.

We also plot the TCP as function of time in Figure 9. In (A) and (C) the virus is injected into the centre of the domain while in (B) and (D) it is injected in five random locations. In (A) and (B) we chose *θ* = 350 and in (C) and (D) we have *θ* = 500. In all cases we see the maximum TCP right after the first cycle. We also note that a distributed injection of virus particles (B) and (D) has a beneficial outcome for the expected tumor control. The plots of the TCP for *θ* = 58.33 are not shown, as the values do not exceed 2 10*^−^*^20^, which we consider to be 0.

**Figure 9:**
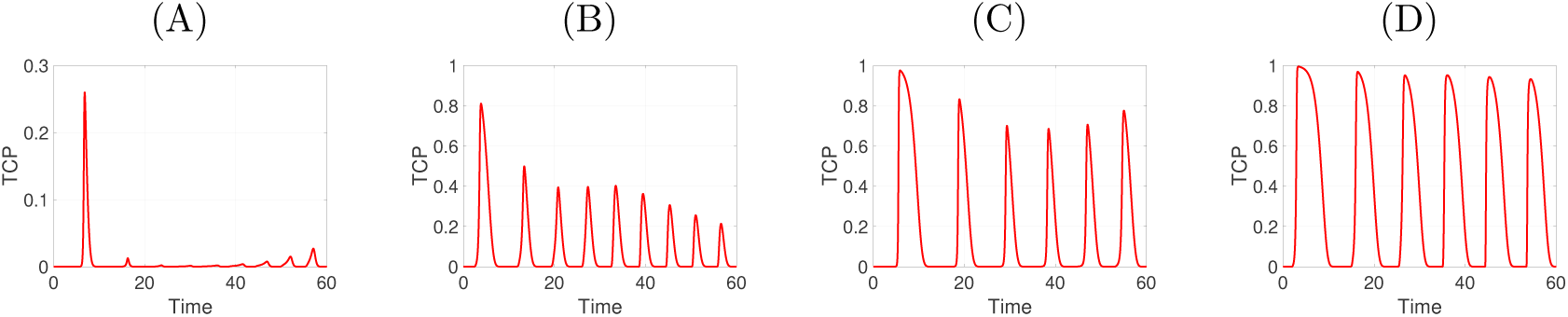
The TCP values of injected tumor cells when *θ > θ_H_* at *a* = 3.33 and *γ* = 13.33. **(A):** *θ* = 350 where the tumor is injected in the center, **(B):** *θ* = 350 where the tumor injected in five random locations, **(C):** *θ* = 500 where the tumor is injected in the center, and **(D):** *θ* = 500 where the tumor is injected five random locations.

We see a significant difference to the TCP values for the ODE and PDE models.

For example for *θ* = 58.33 the TCP in the ODE model was 0.45 while in the spatial model it is 0. Hence the spatial context of cancer and virus spread is very important.

### 4.5. PDE Model 2-D

In this section, we run the simulations of model (3) on a 2-D square domain of dimensions 60 × 60 up to time 37. In dimensional parameters this corresponds to a domain of side length 54 mm and a time of up to 123 days. First, in Figure 10 we choose *θ_t_ < θ < θ_H_* with *θ* = 58.33. Figures (A) and (D) show the cancer cell density with black for high concentration and brown for low. Figures (B) and (E) show the density of infected tumor cells, where high infection is white. Figures (C) and (F) show the distribution of the virus particles from blue (low) to red (high). Figures (A), (B), (C) we show *t* = 10, and Figures (D), (E), (F) are at *t* = 25. The pattern is a filled ring pattern that spreads radially and establishes the coexistence steady state in the inside of the ring.

**Figure 10:**
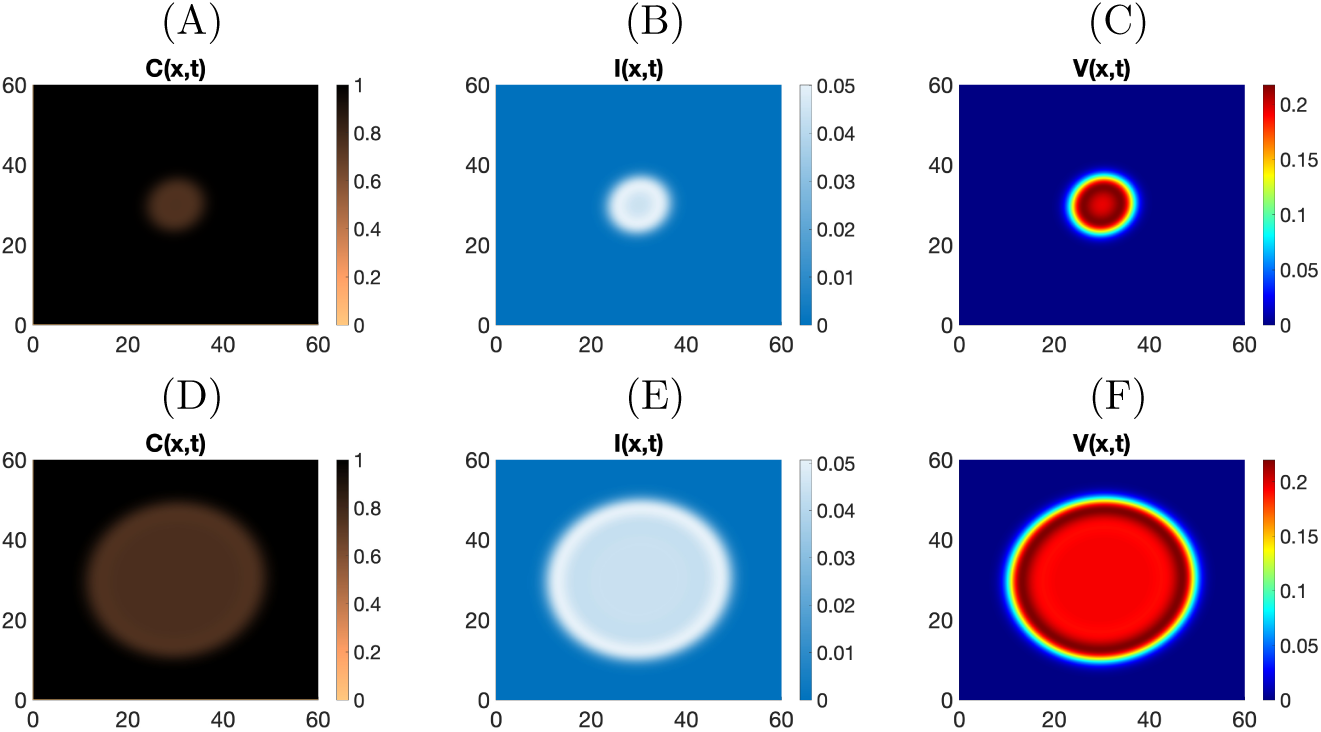
2-D simulations with virus particles injected in the center of the tumor, the spatial distribution of the susceptible cells, infected cells and the virus with *θ* = 58.33, *a* = 3.33 and *γ* = 13.33 at *t* = 10 and *t* = 25, respectively. **(A):** C(x,t), **(B):** I(x,t), and **(C):** V(x,t). **(D):** C(x,t), **(E):** I(x,t), and **(F):** V(x,t).

In Figure 11 we consider the oscillatory range and choose *θ* = 500 *> θ_H_*. We first observe a hollow ring pattern, where tumor is destroyed by the spreading virus wave, leaving a close-to zero state in the inside of the ring. However, as the simulation continues, shown in Figure 12, the cancer and the virus come back. They regrow in the centre of the domain and initiate a second outward moving ring of viral infection. This repeats periodically, though, it is not an exact period, since the inoculation spot in the middle of the domain splits up into two spots as seen in Figure 12 (*t* = 24) and (*t* = 34). This process of periodic spot splitting is a new effect in reaction-diffusion equations and has, as far as we know, never been analysed before.

**Figure 11:**
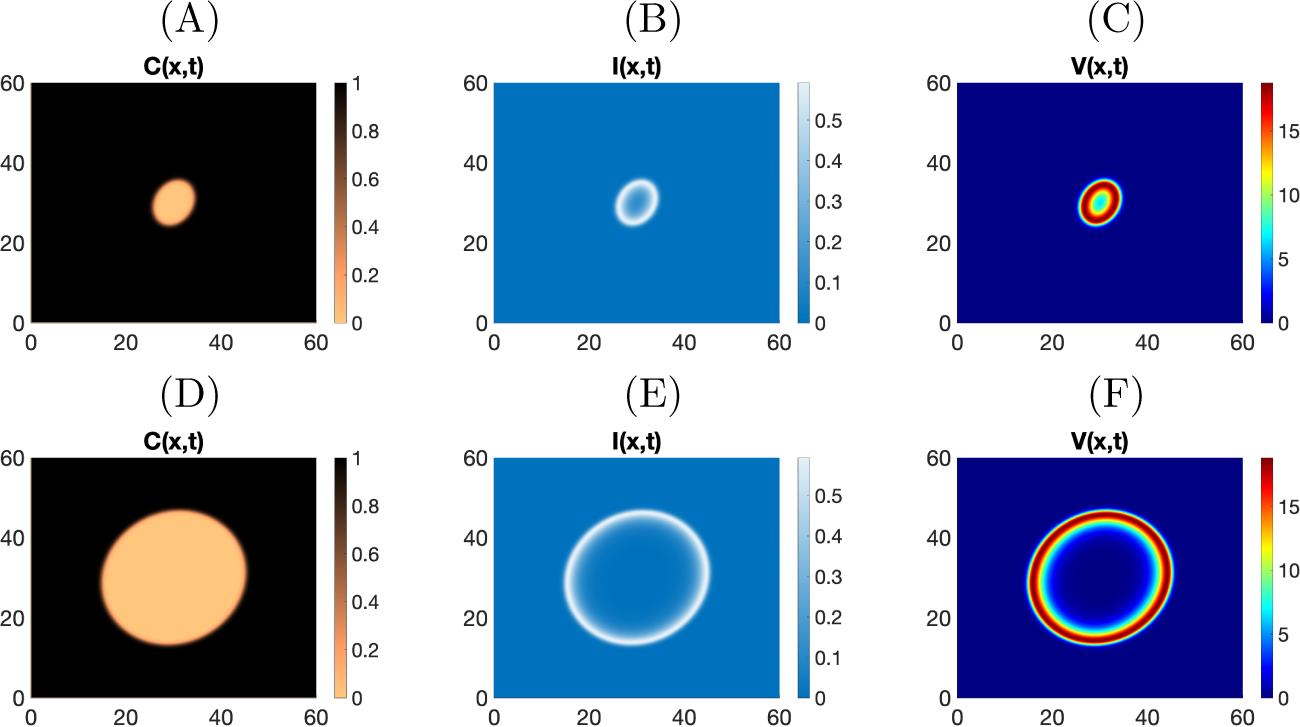
2-D simulations with virus particles injected in the center of the tumor, the spatial distribution of the susceptible cells, infected cells and the virus with *θ* = 500, *a* = 3.33 and *γ* = 13.33 at *t* = 1 and *t* = 3, respectively. **(A):** C(x,t), **(B):** I(x,t), and **(C):** V(x,t). **(D):** C(x,t), **(E):** I(x,t), and **(F):** V(x,t).

**Figure 12:**
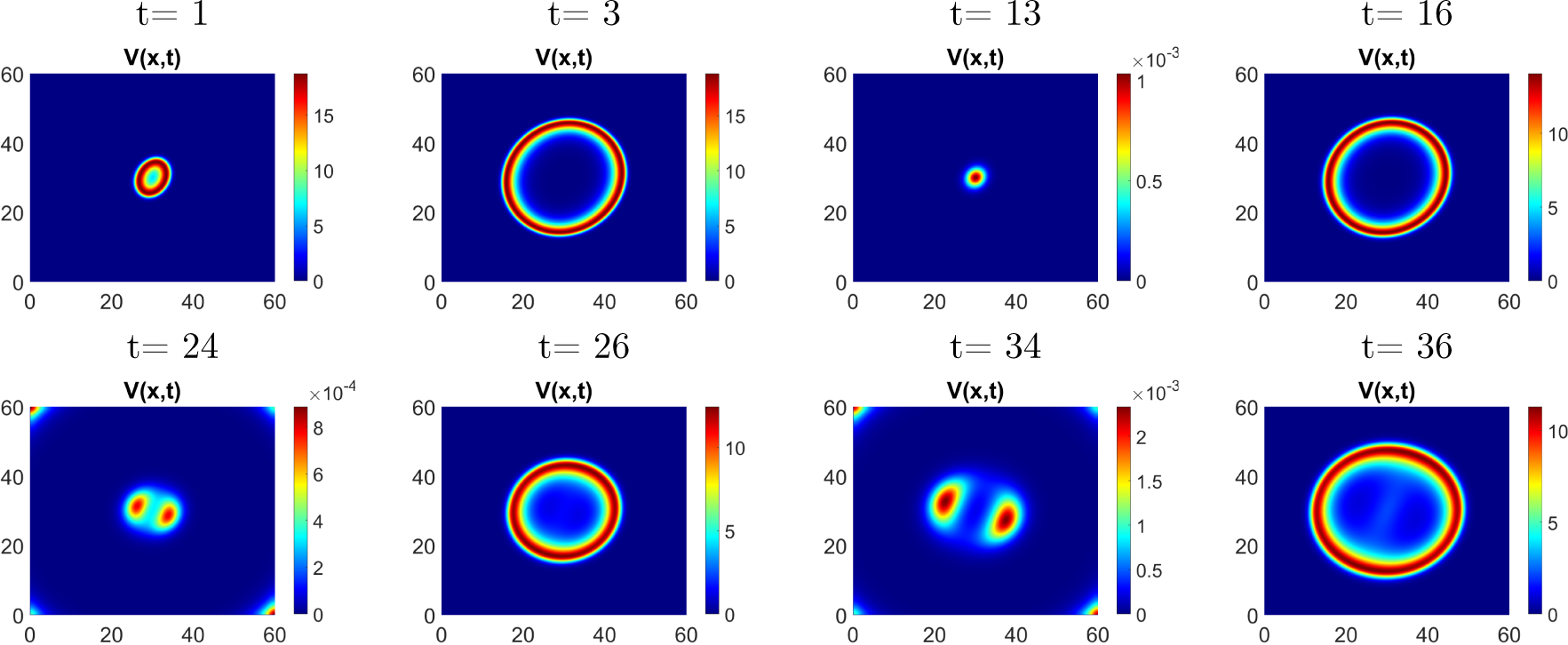
Simulation of model (3) for initial conditions where the virus particles injected in the center of the tumor with *θ* = 500, *a* = 3.33 and *γ* = 13.33.

Similar behaviour has been induced when we use different injection strategies as we can see in Figure 13. The only difference in this case is that the complex interaction of the virus particles and their emergence lead to complex interactions of expanding hollow ring patterns. Again, we observe the periodic peak splitting mentioned above. Finally, we were interested in the long-time dynamics, since it is known that coupled oscillators can lead to chaotic patterns [4; 50; 73]. The MATLAB code used above prevents a very long time analysis, hence we are using a new tool *VisualPDE* that was recently developed at Durham University [88]. This tool allows us to run very long time simulations and also to extend the size of the domain. In Figure 14 we show the long time dynamics of the system on a domain [0, 240]^2^ for the virus component

**Figure 13:**
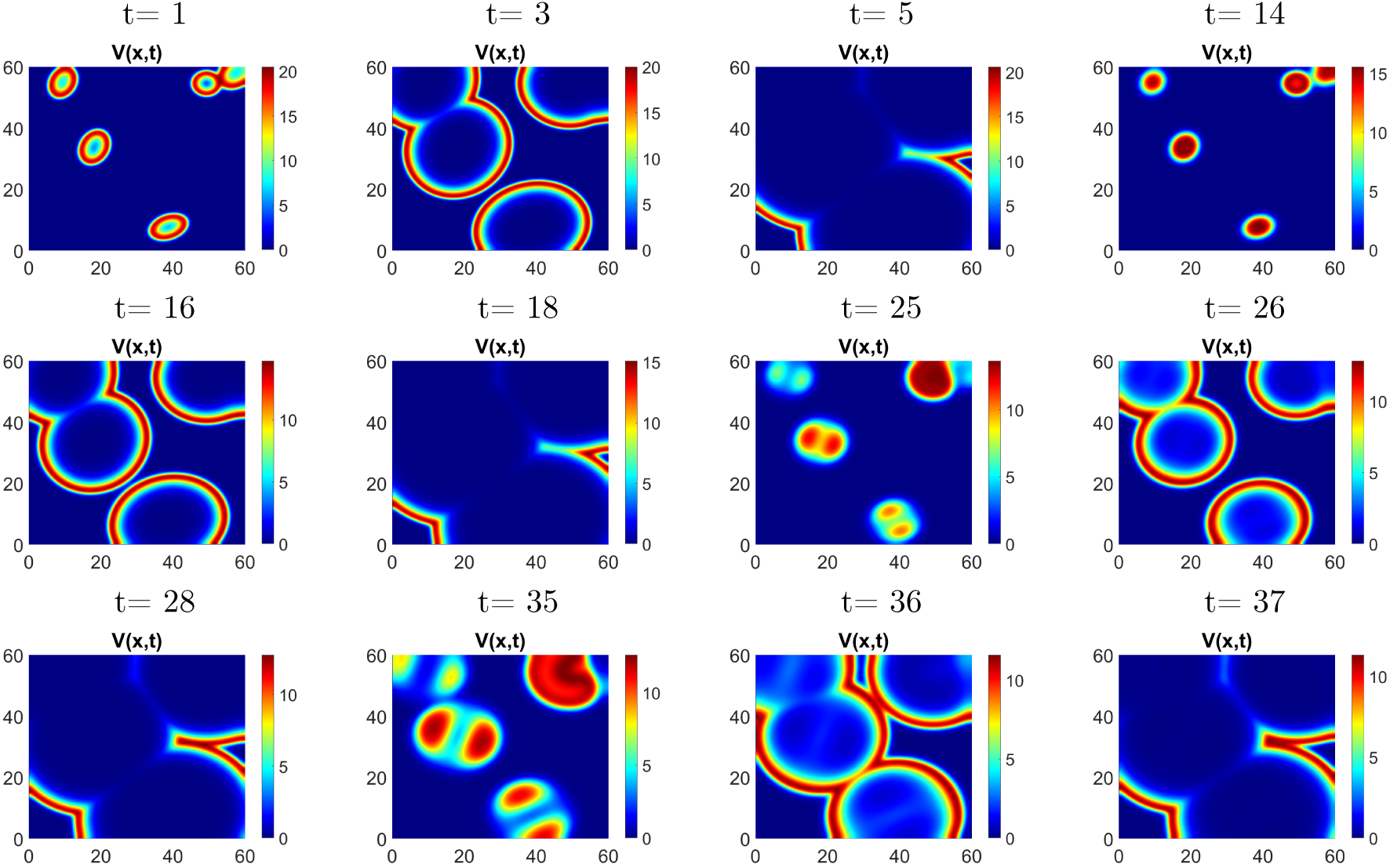
Simulation of model (3) for initial conditions where the virus particles injected in the tumor randomly, with *θ* = 500, *a* = 3.33 and *γ* = 13.33.

**Figure 14:**
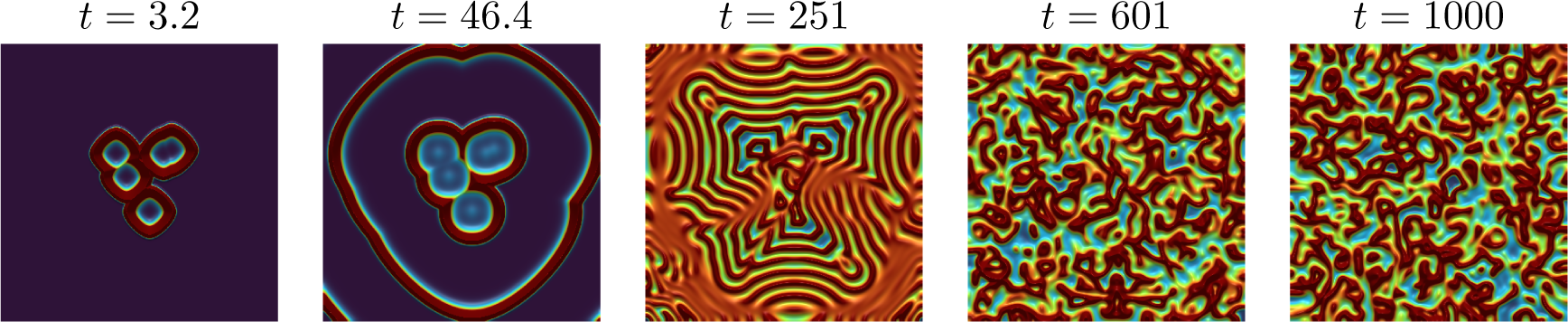
Simulations with *VisualPDE* [88] on a larger domain [0, 240]^2^ with the same initial conditions as used in Figure 13. We show the virus concentration in colors from blue (low) to red (high) with parameters from Table 3 and *θ* = 500.

*V* (*x, t*). We use the same initial conditions as in Figure 13. For short times we see the concentric ring structure, which around time 200-300 creates a complex interaction of spatial periodic solutions. These further interact to form irregular patterns at around *t* = 600. These irregular patterns persist for long times, and a characteristic length scale of these patterns seems to be established (see *t* = 1000). In Figure 15 we show all three components for cancer cells, infected cancer cells, and virus, for the time point *t* = 601. In Figure 16 we consider different values of the bifurcation parameter *θ*. The first choice *θ* = 300 is below the Hopf point, initial oscillations fade out and the solution converges to the homogeneous coexistence steady state. For *θ* = 350 and larger we are past the Hopf point, and oscillations persist, which become more and more irregular for larger values of *θ*.

**Figure 15:**
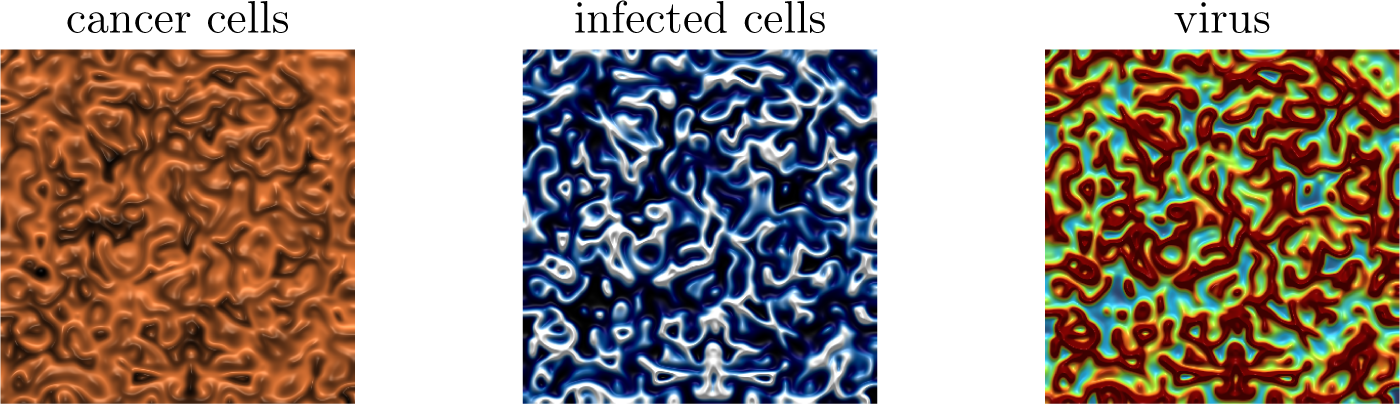
Simulations with *VisualPDE* [88] at time *t* = 601 (2003 days) with parameters from Table 3 and *θ* = 500. The cancer cells on the left are shown from 0 (brown) to 0.2 (black). The scale for the infected cells is 0 (blue) to 0.05 (white), and the scale for the virus is 0 (blue) and 1 (red).

**Figure 16:**
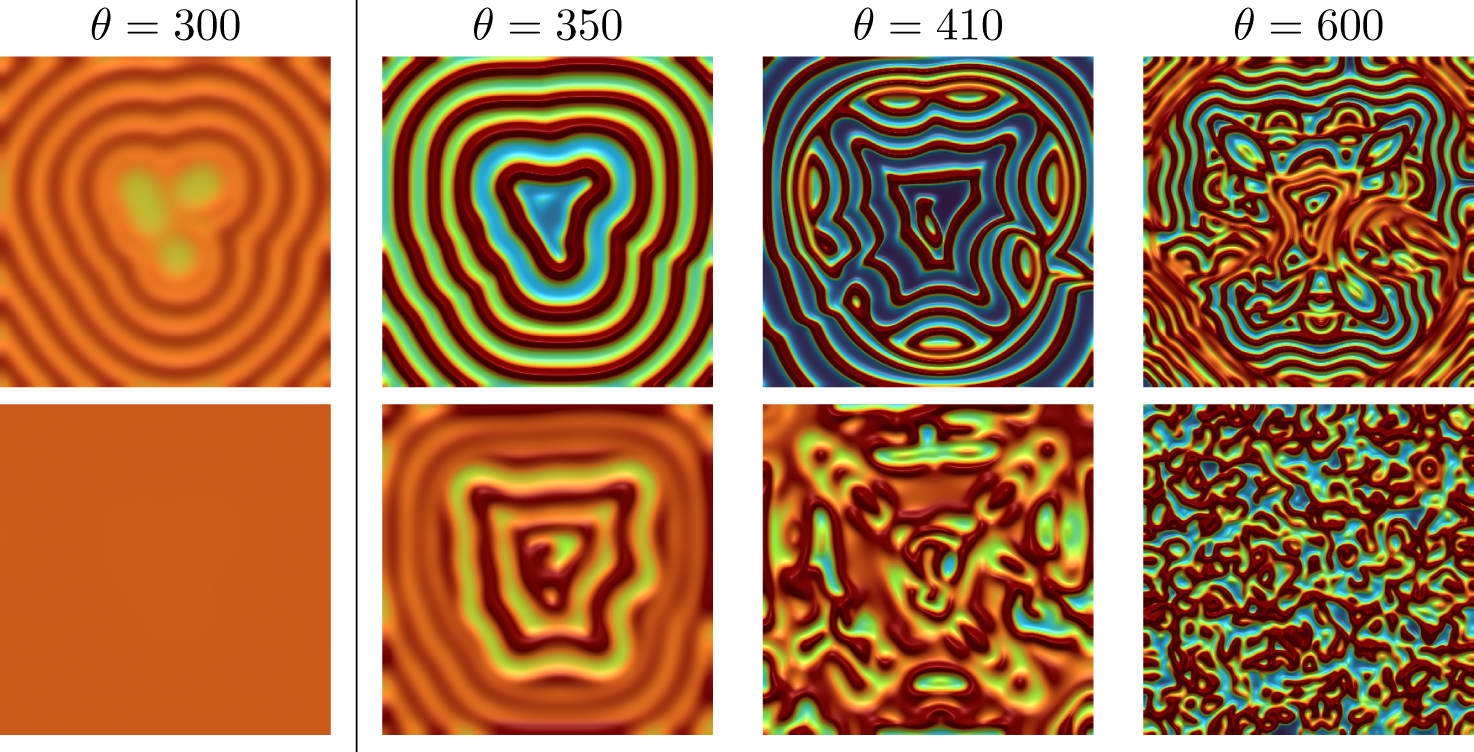
Simulations with *VisualPDE* [88] on the domain [0, 240]^2^ with the same initial conditions as used in Figures 13 and 14. We show the virus concentration in colors from blue (low) to red (high) with parameters from Table 3. The columns represent different values for *θ*. The value of the Hopf bifurcation *θ_H_* = 338.45 is indicated by a vertical line. The rows represent the time points of *t* = 212 and *t* = 601, respectively.

## 5. Conclusion

The mathematical modelling of oncolytic viruses through a susceptible-infected-virus model is a standard approach (see references in the Introduction). The occurance of oscillations is well established and it relates to the predator-prey relationship of the cancer cells and the virus. Two aspects, which were less studied in the literature, are considered here: the relevance of these oscillations in clinical practice and the rich patterns that can arise in the spatial context.

From the medical point of view it is clear that the adaptive immune response is activated within a few days [41]. Hence our simulations beyond a few days are unrealistic for patient outcomes. This includes the prolonged oscillations, which typically had a wave length of 10 days or so. Rather, it appears that the tumor burden goes very close to 0 between these oscillatory outbreaks, indicating tumor removal. We computed the tumor control probability (TCP) to express this fact. We find that with increased viral production rate *θ*, tumor control can be achieved within a few days. However, for realistic values of *θ* = 58.33 we found a TCP of 0 in our spatial model, which confirms the observation that oncolytic viral therapy alone is insufficient in many cases. Mechanisms to increase *θ*seem to be a promising strategy to improve the outcome of oncolytic virotherapy. We recall that

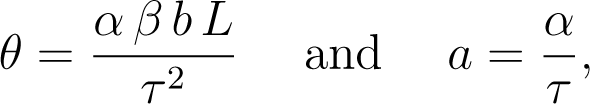

where *α* denotes the removal rate of infected cells, *β* denotes the infection rate, *b* is the burst size of new virions, *L* denotes the carrying capacity of cancer cells and *τ* denotes the cancer growth rate.

An immediate option to increase the efficiency of oncolytic virotherapy is to increase the viral infectivity *β* and the viral burst size *b*. This confirms previous observations by [6; 82; 79] and it is indeed focus of several virology labs to increase the efficiency of a viral infection [12].

The cancer growth dynamics are represented in *θ* as the ratio *L/τ* ^2^. I.e. if the effective growth rate *τ* is large, then *θ* is small and cancer gains an advantage. This opens the door to combine oncolytic virotherapy with growth reduction treatments such as chemotherapy. Chemotherapy can reduce *τ*, thereby moving *θ* into the Hopf region to gain tumor control. Combinations of oncolytic virotherapy and chemotherapy were studied in [85; 14; 63].

The clearance rate of infected cells *α* enters our model parameters in two important ways. On the one hand increasing *α* also increases *θ*. But in addition, increasing *α* increases our bifurcation values for the transcritical bifurcation *θ_t_* and for the Hopf bifurcation *θ_H_*. For the value of *θ_t_* this is obvious from (6). For the Hopf value *θ_H_* we need to look back at the condition *κ*(*θ*) = 0 with *κ*(*θ*) from (8). The coefficient *m* is increasing in *a*, hence all positive terms in *κ*(*θ*) are increasing in *a*. The only negative term is − *θ*^3^. This implies that the positive zero *θ_H_* is also increasing in *a*. As a result, increasing the infected removal rate shifts the system closer to treatment failure. One way how *α* or *a* are increased is the action of the adaptive immune response. If the adaptive immune response activates within a few days, *a* is increased and the dynamic shifts to reduced viral infection and increased cancer growth. This confirms the common understanding that the oncolytic virus has to be fast to accomplish maximal effect before the immune system responds. We did not explicitly model the immune response here, but the effect is clear. Oncolytic virotherapy with inclusion of immune responses was modelled in [1; 23; 79] and others.

Overall we observe a conundrum. The first idea is to control cancer with a virus that is deadly for cancer alone. However, in many cases the virus is not quick enough to spread through the entire tumor before the adaptive immune system kicks in. One strategy to overcome this are multiple virus injections at different sites, as we have shown in our simulations. The second idea is to not expect the oncolytic virus to kill, but rather mark cancer cells with an antigen that is recognized by immune cells. However, immune cell kill might be too quick to allow the viral infection to spread to the entire tumor. A Goldilocks regime needs to be found, where viral infection and immune cell kill balance in the right way. This conclusion is further confirmed by models that include immune response. Storey et al. [79] talk about an intermediate immune response for optimal treatment outcome, and Eftimie et al. [23; 21] show the existence of multi-stability and even multi-instability, which is a strong indication of irregular and chaotic behavior.

In the spatial context we computed the speed of invasion of the virus front. This important information tells us if the virus infection is quick enough to reach the entire tumor, before the adaptive immune system kicks in. In Figure 7 (A), for example, with a single injection in the centre of the tumor, it took about 38 time units (129 days) before the entire domain was infected. A distributed injection in Figure 8 (A) only needed 21 time units (70 days).

In addition, we can do what mathematicians have done ever since modelling existed: forget the biology for the moment, and consider the model as a mathematical object of its own interest. We find that the model reproduces typical viral load patterns such as a coexistence steady state in Figure 10 and a spreading hollow-ring pattern in Figure 11. Moreover, we find a new phenomenon of oscillatory peak splitting (see Figures 12 and 13) which we cannot explain with our current methods. Long time simulations (see Figures 14,15,16) revealed very complex spatio-temporal oscillations, and the analysis of those is currently out of reach.

While these patterns might not be relevant in the context of oncolytic viruses, they can be relevant for other virus infections. For example many COVID-19 patients suffer from long lasting recurring effects ([27],[17]), and we can speculate that continued oscillations might have such an effect. The variability of virus infections is enormous [58], and even our abstract mathematical results might develop into useful tools in the future.

## Acknowledgements

AAB acknowledges support through United Arab Emirates University Scholarship. TH acknowledges support from the Natural Science and Engineering Research Council of Canada (NSERC) RGPIN-2023-04269. AAB and TH thank the members of the Mathematical Biology Journal Club for their valuable comments.

